# A balance index for phylogenetic trees based on quartets

**DOI:** 10.1101/276816

**Authors:** Tomás M. Coronado, Arnau Mir, Francesc Rosselló, Gabriel Valiente

## Abstract

We define a new balance index for phylogenetic trees based on the symmetry of the evolutive history of every quartet of leaves. This index makes sense for polytomic trees and it can be computed in time linear in the number of leaves. We compute its maximum and minimum values for arbitrary and bifurcating trees, and we provide exact formulas for its expected value and variance for bifurcating trees under Ford’s *α*-model and Aldous’ β-model and for arbitrary trees under the α-γ-model.

## 1. Introduction

One of the most broadly studied properties of the topology of phylogenetic trees is their balance, that is, the tendency of the subtrees rooted at all children of any given node to have a similar shape. The main reason for this interest is that the balance of a tree embodies the symmetry of the evolutive history it describes, and hence it reflects, at least to some extent, a feature of the forces that drove the evolution of the set of species considered in the tree [9, Chap. 33].

The symmetry of a tree is usually quantified by means of *balance indices*. Such indices include the *Colless’ index* [7] for bifurcating trees, which is defined as the sum, over all internal nodes *v*, of the absolute value of the difference between the number of descendant leaves of the pair of children of *v*; the *Sackin’s index* [23], which is defined as the sum of the depths of all leaves in the tree, and hence, it measures the average depth of a leaf; and the *total cophenetic index* [17], which is defined as the sum, over all pairs of different leaves of the tree, of the depth of their least common ancestor, and is related to the variance of the leaves’ depths (cf. [17, Lem. 2]). But there are many more balance indices [9, Chap. 33], and Shao and Sokal [24, p. 1990] explicitly advise to use more than one such index to quantify tree balance. Such balance indices only depend on the topology of the trees, not on the branch lengths or the actual taxa labeling their leaves, and they have been widely used as tools to test stochastic models of evolution [3, 12, 13, 18, 24].

In this paper we propose a new balance index, the *quartet index*. It is defined as the sum, over all 4-tuples of different leaves of the tree, of a value that quantifies the symmetry of the joint evolution of the species they represent. The value associated with a 4-tuple of leaves is simply required to increase with the number of isomorphisms of the phylogenetic subtree (the *quartet)* they induce. The main features of our index are that it can be easily computed in linear time and that its expected value and variance can be explicitly computed on any probabilistic model of phylogenetic trees satisfying two natural conditions: independence under relabelings and sampling consistency. This allows us to provide these values for two well-known probabilistic models for bifurcating phylogenetic trees, Ford’s α-model [10] and Aldous’ β-model [1], which include as specific instances the Yule *[11, 27]* and the uniform *[5, 22, 26]* models, as well as for Chen-Ford*’s α*-γ-model for polytomic trees [6]. To our knowledge, this is the first shape index for which formulas for the expected value and the variance under the α-γ-model have been provided. We also compute the maximum and minimum values of our quartet index, in both the arbitrary and the bifurcating cases: the minimum value is reached exactly at the combs and the maximum value is reached exactly at the rooted stars in the arbitrary case and at the maximally balanced trees in the bifurcating case.

The rest of this paper is organized as follows. In a first section we introduce the basic notations and facts on phylogenetic trees that will be used in the rest of the paper, and we recall several preliminary results on probabilistic models of phylogenetic trees, proving those results for which we have not been able to find a suitable reference in the literature. Then, in Section 3, we define our quartet index *QI* and we establish its basic properties. In Section 4 we compute its maximum and minimum values, and finally, in Section 5, we compute its expected value and variance under different probabilistic models. This paper is accompanied by the GitHub page https://github.com/biocom-uib/Quartet_Index containing a set of Python scripts that perform several computations related to this index.

## 2. Preliminaries

### 2.1 Notations and conventions

In this paper, by an (*unlabeled) tree* we mean a rooted tree without out-degree 1 nodes. As it is usual, we understand such a tree as a directed graph, with its arcs pointing away from the root. A tree is *bifurcating* when all its internal nodes have out-degree 2. We shall denote by *L*(*T)* the set of leaves of a tree *T,* by *V*_*int*_(*T)* its set of internal nodes, and by child(*u*) the set of *children* of an internal node *u*, that is, those nodes *v* such that (*u, v*) is an arc in *T*. We shall always consider two isomorphic trees as equal, and we shall denote by 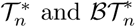 the sets of (isomorphism classes of) trees and of bifurcating trees with *n* leaves, respectively.

A *phylogenetic tree* on a set ∑ is a tree with its leaves bijectively labeled in ∑. An *isomorphism* of phylogenetic trees is an isomorphism of trees that preserves the leaves’ labels. To simplify the language, we shall always identify a leaf of a phylogenetic tree with its label and we shall say that two isomorphic phylogenetic trees “are the same”. We shall denote by *𝒯* (∑) and ℬ𝒯 (∑) the sets of (isomorphism classes of) phylogenetic trees and of bifurcating phylogenetic trees on ∑, respectively. If ∑ and ∑^*′*^ are any two sets of labels of the same cardinal, say *n*, then any bijection ∑↔ ∑^*′*^ extends in a natural way to bijections 𝒯 (∑) ↔ (∑^*′*^) and ℬ𝒯 (∑) ↔ℬ𝒯 (∑^*′*^). When the specific set of labels ∑ is unrelevant and only its cardinal matters, we shall write 𝒯_*n*_ and ℬ𝒯_*n*_ (with *n* = |∑|) instead of 𝒯(∑) and ℬ𝒯 (∑), and we shall identify ∑ with the set [*n*] = {1, 2,*…, n}.* If |∑| = *n*, there exists a forgetful mapping 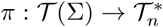 that sends every phylogenetic tree *T* on ∑ to its underlying unlabeled tree: we shall call *π*(𝒯) the *shape* of *T.* We shall write *T*_1_≡*T*_2_ to denote that two phylogenetic trees *T*_1_, *T*_2_ (possibly on different sets of labels of the same cardinal) have the same shape.

We shall represent trees and phylogenetic trees by means of their usual Newick format [19], although we shall omit the ending mark “;” in order not to confuse it in the text with a semicolon punctuation mark. In the case of trees, we shall denote the leaves with symbols *.

Given two nodes *u, v* in a tree *T,* we say that *v* is a *descendant* of *u*, and also that *u* is an *ancestor* of *v*, when there exists a path from *u* to *v* in *T*; this, of course, includes the case of the stationary path from a node *u* to itself, and hence, in this context, we shall use the adjective *proper* to mean that *u ≠ v*. Given a node *v* of a tree *T,* the *subtree T*_*v*_ *of T rooted at v* is the subgraph of *T* induced by the descendants of *v*. We shall denote by *κ*_*T*_ (*v*), or simply by *κ*(*v*) if *T* is implicitly understood, the number of leaves of *T*_*v*_.

Given a tree *T* and a subset *X*⊆*L*(*T)*, the *restriction T* (*X*) of *T* to *X* is the tree obtained by first taking the subgraph of *T* induced by all the ancestors of leaves in *X* and then suppressing its out-degree 1 nodes. By *suppressing* a node *u* with out-degree 1 we mean that if *u* is the root, we remove it together with the arc incident to it, and, if *u* is not the root and if *u′* and *u″* are, respectively, its parent and its child, then we remove the node *u* and the arcs (*u′, u*), (*u, u″*) and we replace them by a new arc (*u′, u″*). For every *Y*⊆*L*(*T)*, the tree *T* (-*Y)* obtained by *removing Y* from *T* is nothing but the restriction *T* (*L*(*T)* \*Y)*. If *T* is phylogenetic tree on a set Σ and *X*⊆ *∑*, the restrictions *T* (*X*) and *T* (-*X*) are phylogenetic trees on *X* and ∑\*X*, respectively.

A *comb* is a bifurcating phylogenetic tree such that all its internal nodes have a leaf child: see Fig. 1.(a). All combs with the same number *n* of leaves have the same shape, and we shall generically denote them (as well as their shape in 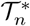) by *K*_n_. A *star* is a phylogenetic tree all whose leaves are children of the root: see Fig. 1.(b). For every set ∑, there is only one star on ∑, and if *|∑|* = *n*, we shall generically denote it (as well as its shape) by *S*_n_.

**Figure 1:**
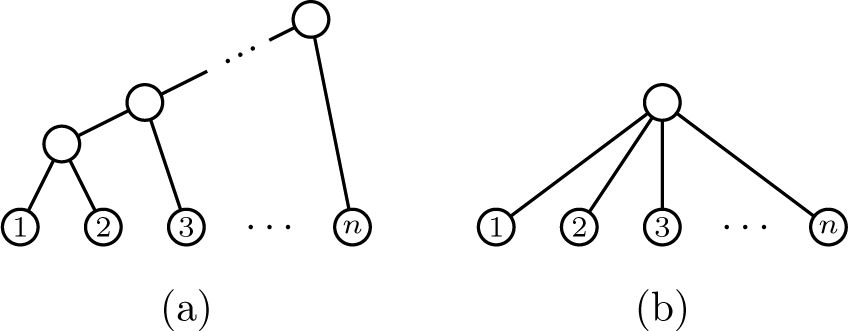
(a) A comb *K*_*n*_. (b) A star *S*_*n*_.

Given *k⩾* 2 phylogenetic trees *T*_1_,*…, T*_*k*_, with every *T*_*i.*_ ∈ 𝒯 (∑_*i*_.) and the sets of labels ∑_*i*_. pairwise disjoint, their *root join* is the phylogenetic tree *T*_*1*_*⋆T*_2_⋆_…._⋆T_*k*_ on 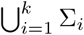 obtained by connecting the roots of (disjoint copies of) *T*_1_,*…, T*_*k*_ to a new common root *r*; see Fig. 2. If *T*_1_,*…, T*_*k*_ are unlabeled trees, a similar construction yields a tree *T*_1_ ⋆…⋆*T*_*k*_.

**Figure 2:**
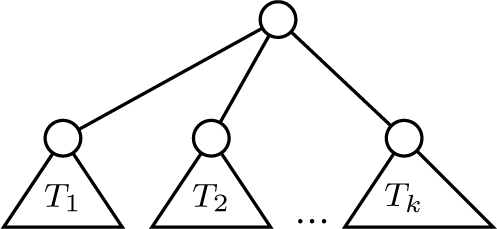
The root join *T*_1_⋆…⋆*T*_*k*_.

Let *T* be a bifurcating tree. For every *v ∈V*_*int*_(*T)*, say with children *v*_1_, *v*_2_, the *balance value* of *v* is *bal*_*T*_ (*v*) = │*κ*(*v*_1_)-*κ*(*v*_2_)│. An internal node *v* of *T* is *balanced* when *bal*_*T*_ (*v*) ⩽ 1. So, a node *v* with children *v*_1_ and *v*_2_ is balanced if, and only if, {*κ*(*v*_1_), *κ*(*v*_2_)} = {⌊*κ*(*v*)*/*2⌋, ⌊*κ*(*v*)*/*2⌋}. We shall say that a bifurcating tree *T* is *maximally balanced* when all its internal nodes are balanced. Recurrently, a bifurcating tree is maximally balanced when its root is balanced and the subtrees rooted at the children of the root are both maximally balanced. This implies that, for any fixed number *n* of nodes, there is only one maximally balanced tree in 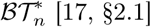

### 2.2 Probabilistic models

A *probabilistic model of phylogenetic trees* (*P*_*n*_)_*n*_ is a family of probability mappings *P*_n_ : т_n_ *⟶* [0, 1], each one sending each phylogenetic tree in т_n_ to its probability under this model. Every such a probabilistic model of phylogenetic trees (*P*_n_)_n_ induces a *probabilistic model of trees*, that is, a family of probability mappings 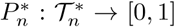, by defining the probability of a tree as the sum of the probabilities of all phylogenetic trees in Τ_n_ with that shape:

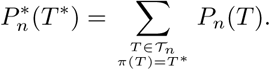

If *|∑|* = *n*, then *P*_n_ : т_n_ *⟶* [0, 1] induces also a probability mapping *P∑* on т (∑) through the bijection *T∑* _⟷_ т_n_ induced by a given bijection ∑ ⟷ [*n*].

*A probabilistic model of bifurcating phylogenetic trees* is a probabilistic model of phylogenetic trees (*P*_n_)_n_ such that *P*_n_(т) = 0 for every *T ∈𝒯*_n.\_ *ℬ𝒯*_n_

A probabilistic model of phylogenetic trees (*P*_n_)_n_ is *shape invariant* (or *exchangeable* [1]) when, for every T, T ^*′*^ *∈ 𝒯*_n_, if *T = T* ^*′*^, then *P*_n_(*T)* = *P*_n_(*T* ^*′*^). In this case, for every 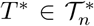 and for every *T ∈ π*^*-*1^(*T* ^*^),

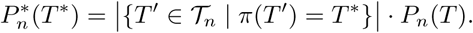

Conversely, every probabilistic model of trees 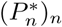 defines a shape invariant probabilistic model of phylogenetic trees (*P*_n_)_n_ by means of

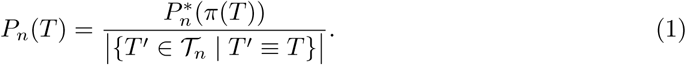

Notice that if (*P*_*n*_)_*n*_ is shape invariant, then, for every set of labels Σ, say, with *|∑|* = *n*, the probability mapping *P∑* : ℐ (∑) ⟶ [0, 1] induced by (*P*_*n*_)_*n*_ does not depend on the specific bijection *∑* ↔ [*n*] used to define it.

A probabilistic model of phylogenetic trees (*P*_*n*_)_*n*_ is *sampling consistent* [1] *(or also deletion stable* [10]) when, for every *n≥* 2, if we choose a tree *T ∈ 𝒯*_*n*_ with probability distribution *P*_*n*_ and we remove its leaf *n*, the resulting tree is obtained with probability *P*_*n-*1_; formally, when, for every *n≥* 2 and for every *T*_0_ ∈ *𝒯*_n-1_,

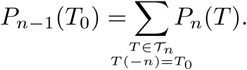

It is straightforward to prove, by induction on *n − m* and using that, for every *T ∈𝒯* _*n*_ and for every 1≤ *m < n*, the restriction of *T* (-*n*) to [*m*] is simply *T* ([*m*]), that this condition is equivalent to the following: (*P*_*n*_)_*n*_ is sampling consistent when, for every *n≥* 2, for every 1 ≤*m < n*, and for every *T*_0_ ∈ *𝒯*_*m*_,

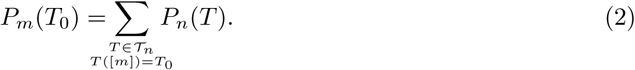

It is also easy to prove that if (*P*_n_)_n_ is sampling consistent *and* shape invariant, so that the probability of a phylogenetic tree is not affected by permutations of its leaves, then, for every *n ≥* 2, for every ⊘≠⊊; [*n*], say, with *|X|* = *m*, and for every *T*_0_ ∈ *𝒯* (*X*),

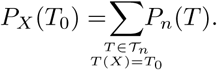

(where *P*_X_ stands for the probability mapping on 𝒯 (*X*) induced by *P*_m_ through any bijection *X ↔ ([m])*.

A probabilistic model of trees 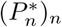 is *sampling consistent*, or *deletion stable*, when, for every *n ≥* 2, if we choose a tree 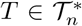 with probability distribution 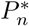 and a leaf *x ∈ L*(*T)* equiprobably and if we remove *x* from *T,* the resulting tree is obtained with probability 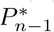:formally, when, for every *n≥ 2* and for every 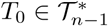,

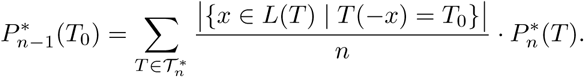

We prove now several lemmas on probabilistic models that will be used in §5. The first lemma provides an extension of equation (2) to trees; we include it because we have not been able to find a suitable reference for it in the literature. In it, and henceforth, 𝒫_k_(*X*) denotes the set of all subsets of cardinal *k* of *X*.

**Lemma 1.** *A probabilistic model of trees* 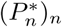 *is sampling consistent if, and only if, for every n* ≥ 2, *for every* 1≤*m < n, and for every* 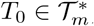,

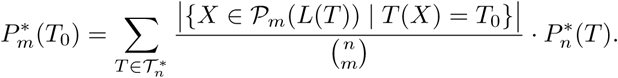

*Proof.* The “if” implication is obvious. As far as the “only if” implication, we prove by induction on *n - m* that if 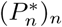 (is sampling consistent, then, for every 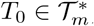,

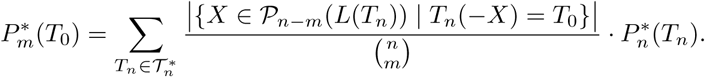

The starting case *m* = *n-*1 is the sampling consistency property. Assume now that this equality holds for *m* and let 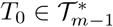. Then

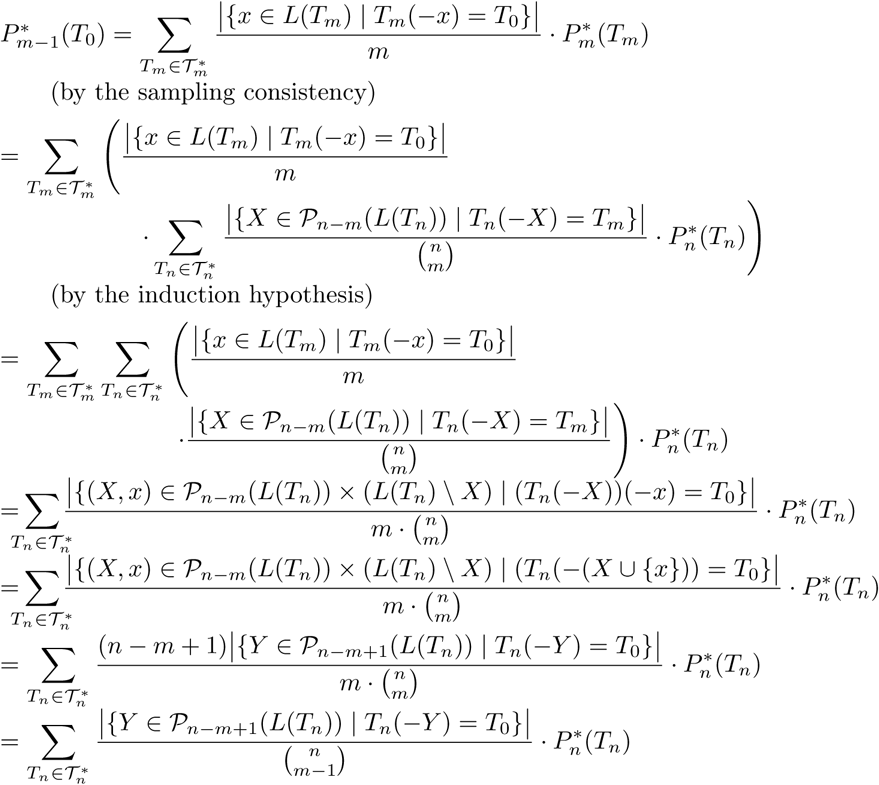

which proves the inductive step.

**Lemma 2.** *Let* (*P*_*n*_)_*n*_ *be a shape invariant probabilistic model of phylogenetic trees. For every* 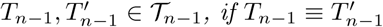 *then*

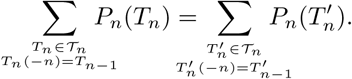

*Proof.* Let 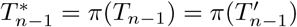 and let 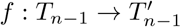 be an isomorphism of unlabeled trees, which exists because *T*_*n-*1_ and 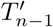 are both isomorphic as unlabeled trees to their shape 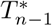 For every *T ∈ 𝒯*_*n-*1_, let

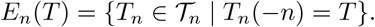

Each *T*_n_ in *E*_n_(*T*_n*-*1_) is obtained by adding a leaf *n* to *T*_*n-*1_ as a new child either to an internal node, or to a new node obtained by splitting an arc into two consecutive arcs, or to a new bifurcating root (whose other child will be the old root). This entails the existence of a shape preserving bijection

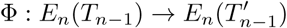

that sends each *T*_n_ *∈ E*_n_(*T*_*n-1*_) to the phylogenetic tree Ф (*T*_n_) obtained by adding the leaf *n* to 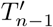 at the place corresponding through the isomorphism *f* to the place where it has been added to *T*_*n-*1_. Then, since (*P*_n_)_n_ is shape invariant,

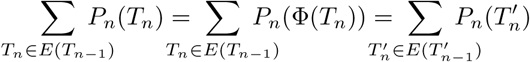

as we claimed.

Next lemma generalizes Cor. 40 in [10]. For the sake of completeness, we provide a direct complete proof of it.

**Lemma 3.** *Let* (*P*_n_)_n_ *be a shape invariant probabilistic model of phylogenetic trees and let* 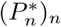 *be the corresponding probabilistic model of trees. Then,* (*P*_n_)_n_ *is sampling consistent if, and only if,* 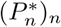 *is sampling consistent.*

*Proof.* Let us prove first the “only if” implication. Let (*P*_n_)_n_ be sampling consistent. Then, for every 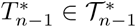 and for every 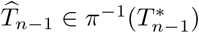,

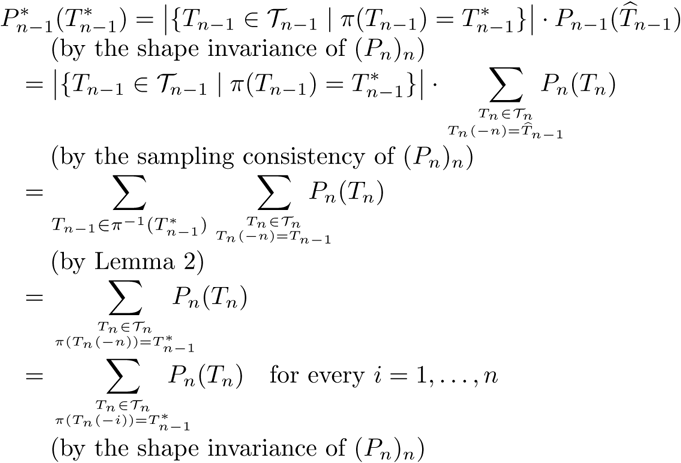

Therefore

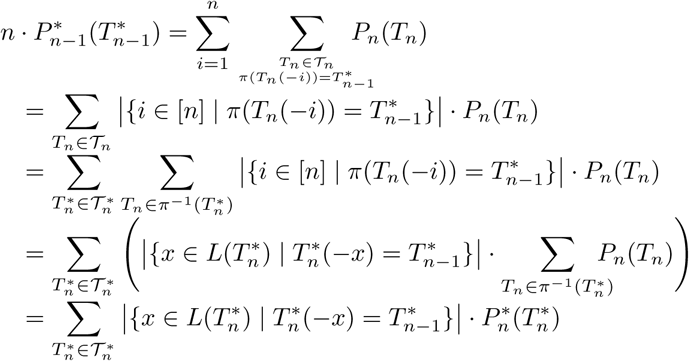

and hence

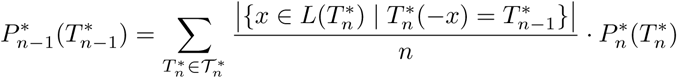

as we wanted to prove.

The proof on the “if” implication consists in carefully running backwards the sequence of equalities in the proof of the “only if” implication. Indeed, assume that 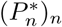 is sampling consistent and let *T*_*n-*1_ ∈ *𝒯*_*n-1*_ and 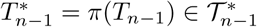. Then

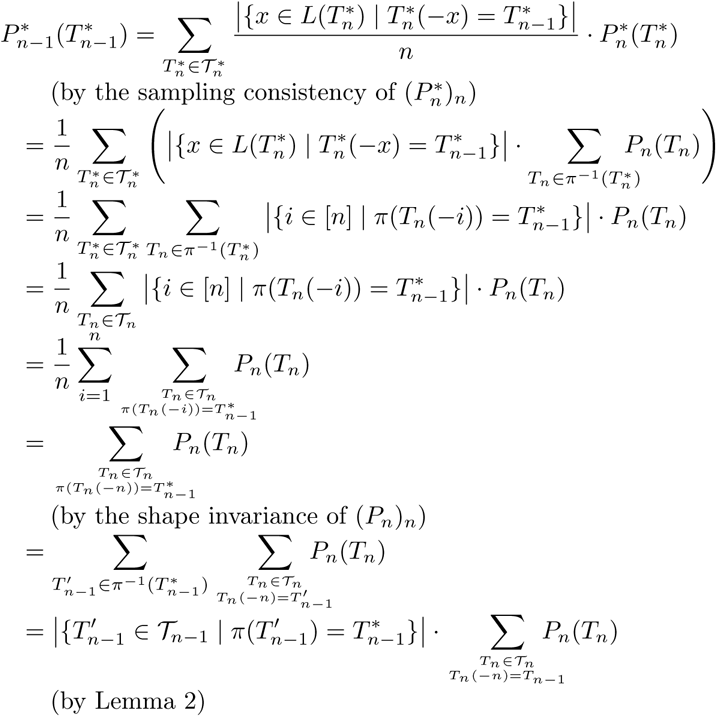

and thus, dividing both sides of this equality by 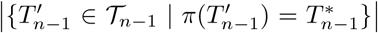 and using the shape invariance of (*P*_n_)_n_, we obtain

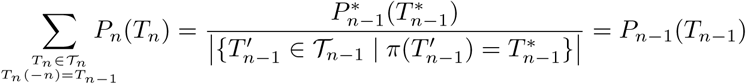

as we wanted to prove.

In §5 we shall be concerned with three specific parametric probabilistic models of phylogenetic trees:

*1)Aldous’ β-model*. The β-splitting model 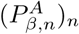 [1, 2] is a probabilistic model *of* bifurcating phylogenetic trees that depends on one parameter β ∈] − 2, *∞* [. Under it, the probability 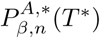 of a tree *T\** with *n* leaves is computed through all possible ways of building it from a single node by recurrently splitting leaves into cherries with leaves having two predetermined target numbers of descendant leaves in the final tree; the parameter β controls the probabilities of these target numbers of descendant leaves. Then, the probability 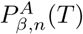of a phylogenetic tree *T* is obtained from the probability of its shape *T\** by means of equation (1). In this way, 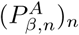 is shape invariant by construction, and it turns out to be sampling consistent [1, §6.3];hence, by Lemma 3, the β-model of trees 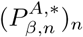 is also sampling consistent. The β-model includes as specific cases the Yule model [11, 27] (when β = 0) and the uniform model [5, 21] (whenβ = −3*/*2).

In §5 we shall need to know the probability 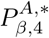 of the maximally balanced tree with 4 leaves ((*\*, \**), (*\*, \**)), which we denote in this paper by 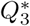 (see Figure 3). This probability is

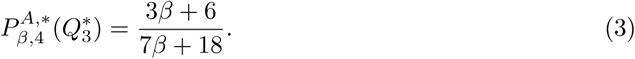

For the convenience of the reader, we compute it in the Appendix A.1 at the end of the paper (see Lemma 25).

*2)Ford’s α-model.* The α-model 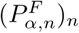 [10] is another probabilistic model of bifurcating phylogenetic trees that depends on one parameter α ∈ [0, 1]. In it, the probability 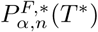 of a tree *T\** is determined through all possible ways of constructing a phylogenetic tree with this shape from a single node (labeled with 1) by adding recurrently new leaves labeled with 2,*…, n* within arcs or to a new root, with the parameter *α* controlling the probability of adding the new leaf within an arc ending in a leaf or elsewhere. Then, the probability 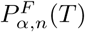 of a phylogenetic tree *T* is obtained from the probability of its shape by means of equation (1). The *α*-model is shape invariant by construction and sampling consistent by [10, Prop. 42], and it also includes as specific cases the Yule model (when α = 0) and the uniform model (when α = 1*/*2).

In §5, we shall also need to know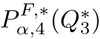, which is provided by Ford in [10, §7: Fig. 20]:

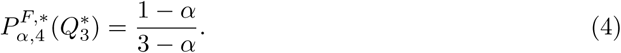

*3)Chen-Ford-Winkel’s α-γ-model*. The *α*-*γ*-model (*P*_α,γ,n_)_n_ [6] *is a probabilistic model of phylogenetic trees that depends on two parameters 0 ≤ γ ≤ α ≤* 1.*≤* In it, the probability of a phylogenetic tree is computed as it is built from a single node (labeled with 1) by adding recurrently new leaves labeled with 2,*…, n* to a new root, to internal nodes, or within arcs, with the parameters α, *γ* controlling the probabilities of choosing the place where to add a new leaf. It turns out that (*P*_α,γ,n_)_n_ is not shape invariant in general [6, Prop. 1.(b)], but that the corresponding model for trees 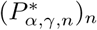 is sampling consistent [6, Thm. 2]. When α = γ, 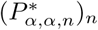 is simply the *α*-model 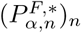 for unlabeled trees.

**Figure 3:**
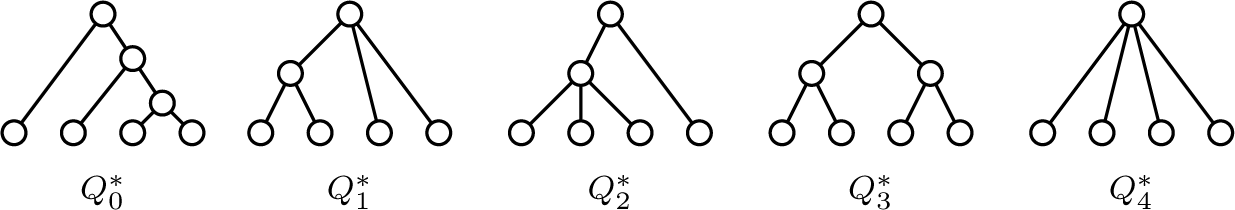
The 5 trees in 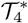

Later in this paper we shall need to know the probabilities under 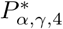 of the five different trees in 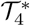, described in Figure 3. They are:

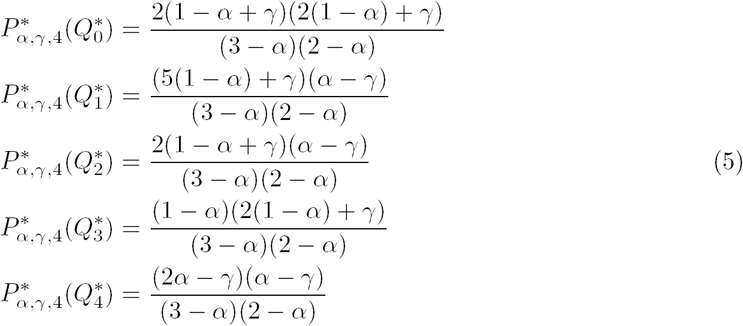

For the convenience of the reader, we also compute these probabilities in the Appendix A.2 at the end of the paper (see Lemma 26).

## 3. Quartet indices

Let *T* be a phylogenetic tree on a set Σ. For every *Q∈𝒫*_4_(Σ), the *quartet on Q displayed* by *T* is the restriction *T* (*Q*) of *T* to *Q*. A phylogenetic tree *T∈𝒯*_*n*_ can contain quartets of five different shapes; they are listed in Figure 3, together with the notations used in this paper to denote them (motivated by Table 1). Notice that a bifurcating phylogenetic tree *T∈ℬ𝒯*_n_ can only contain quartets of two shapes: those denoted by 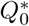 and 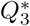 in the aforementioned figure.

We associate to each quartet a *QI-value q*_i_ that increases with the symmetry of the quartet’s shape, as measured by means of its number of automorphisms, going from a value *q*_0_ = 0 for the least symmetric tree, the comb 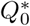, to a largest value of *q*_4_ for the most symmetric one, the star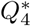; see Table 1. The specific numerical values can be chosen in order to magnify the differences in symmetry between specific pairs of trees. For instance, one could take *q*_*i*_ = *i*, or *q*_*i*_ = 2^*i*^.

**Table 1:**
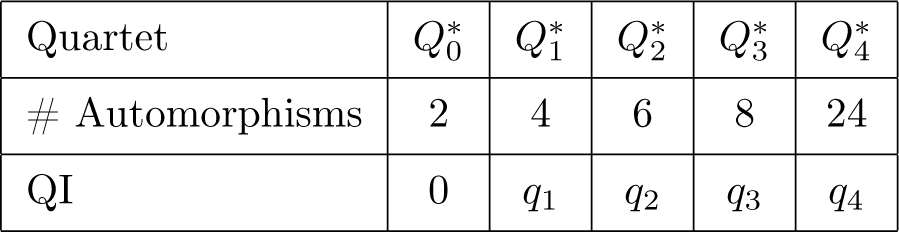
The quartets’ QI-values, with 0 *< q*_1_ *< q*_2_ *< q*_3_ *< q*_4_.

| Quartet | $Q_0^*$ | $Q_1^*$ | $Q_2^*$ | $Q_3^*$ | $Q_4^*$ |
| --- | --- | --- | --- | --- | --- |
| # Automorphisms | 2 | 4 | 6 | 8 | 24 |
| QI | 0 | $q_1$ | $q_2$ | $q_3$ | $q_4$ |

Now, for every *T ∈ 𝒯* (∑), we define its *quartet index QI*(*T)* as the sum of the *QI*-values of its quartets:

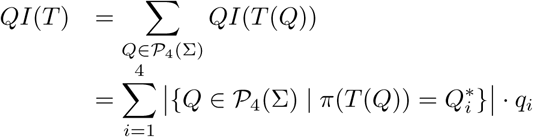

In particular, if |Σ|≤ 3, then *QI*(*T)* = 0 for every *T ∈𝒯* (∑). So, *we shall assume henceforth* that|∑ |≥4.

It is clear that *QI* is a *shape index*, in the sense that two phylogenetic trees with the same shape have the same quartet index. It makes sense then to define the quartet index *QI*(*T\**) of a tree 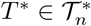 as the quartet index of any phylogenetic tree of shape *T\**.

**Example 4.** Consider the tree *T* = ((1, 2, 3), 4, (5, (6, 7))) depicted in Figure 4. It has: 4 quartets of shape 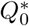; 18 quartets of shape 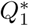; 4 quartets of shape 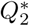; 9 quartets of shape 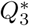; and no quartet of shape 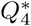 Therefore

**Figure 4:**
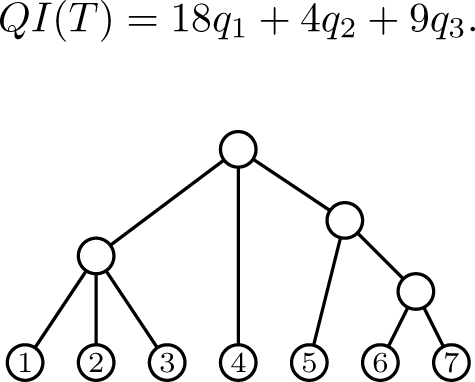
The tree ((1, 2, 3), 4, (5, (6, 7))).

**Remark 5.** If we do not take *q*_0_ = 0, then the resulting index would be

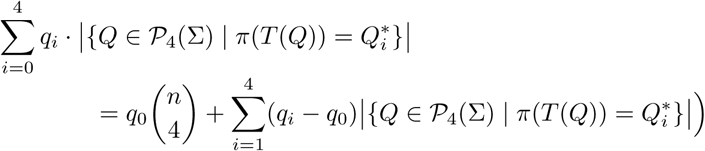

which is equivalent (up to the constant addend 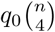) to *QI* taking as *QI*-values 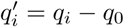

**Remark 6.** One could also associate other values to the quartet shapes; for instance their Sackin index [23, 24] or their total cophenetic index [17], which measure the unbalance of the quartet’s shape, from a smallest value at 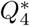 to a largest value at 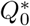. All results obtained in this paper are easily translated to any other sets of values.

Since a bifurcating tree can only contain quartets of shape 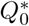 and 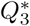, its *QI* index is simply *q*_3_ times its number of quartets of shape 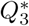. Therefore, in order to avoid this spurious factor, when dealing only with bifurcating trees we shall use the following alternative *quartet index for bifurcating trees QIB*: for every *T ℬ𝒯*(∑),

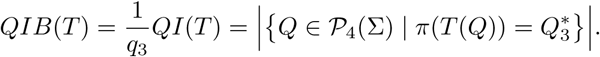

The quartet index for bifurcating trees satisfies the following recurrence.

**Lemma 7.** *Let T* = *T*_1_ *\* T*_2_ ∈ *ℬ𝒯*_*n*_, *where each T*_*i*_ *has n*_*i*_*leaves. Then,*

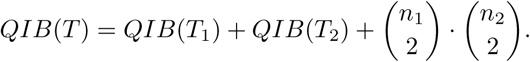

*Proof.* For every *Q ∈ 𝒫*_4_([*n*]), there are the following possibilities:

1. If *Q* ⊆ *L*(*T*_*i*_.), for some *i* = 1, 2, then *T* (*Q*) = *T*_*i*_.*Q*). Therefore, the quartets with *Q ⊆ L*(*T*_*i*_.) contribute *QIB*(*T*_*i*_) to *QIB*(*T)*.
2. If three leaves in *Q* belong to one of the subtrees *T*_*i*_and the fourth to the other subtree *T*_*j*_, then *T* (*Q*) has shape 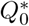 and thus it does not contribute anything to *QIB*(*T)*.
3. If two leaves in *Q* belong to *T*_1_ and the other two to *T*_2_, then *T* (*Q*) has shape 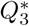 and thus it contributes 1 to *QIB*(*T)*. There are 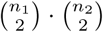 such quartets.

Thus, *QIB* is a *bifurcating recursive tree shape statistic* in the sense of [16]. The recurrence in the last lemma implies directly the following explicit formula for *QIB*, which in particular entails that it can be easily computed in time *O*(*n*), with *n* the number of leaves of the tree, by traversing the tree in post-order (cf. the first paragraph in the proof of Proposition 10 below):

**Corollary 8.** *If, for every T ∈ ℬ𝒯*_*n*_*and for every v ∈ V*_*int*_(*T), we set* child(*v*) = *{v*_1_, *v*_2_*}, then*

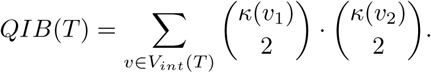

Unfortunately, *QI* is not recursive in this sense: there does not exist any family of mappings (*q*_*m*_>:ℕ^*m*^*→ ℝ*)_*m≥*2_ such that, for every *T∈𝒯* _*n*_, if *T* = *T*_1_ *\* … * T*_*m*_, with each *T*_*i*_having *n*_*i*_leaves, then

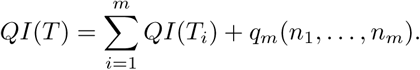

However, next lemma shows that there exists a slightly more involved linear recurrence for *QI*, with its independent term depending on more indices of the trees *T*_*i*_than only their numbers of leaves, which still allows its computation in linear time.

For every *T ∈ 𝒯*_*n*_, let ?(*T*) be the number of *triples* in *T* (that is, of restrictions of *T* to sets of 3 leaves) of shape *S*_3_. Notice that if *T* = *T*_1_ *\* · · · * T*_*m*_and *|L*(*T*_*i*_.)*|* = *n*_*i*_, for each *i* = 1,*…, m*, then

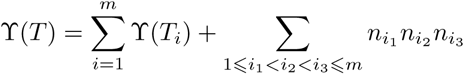

and hence

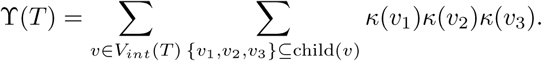

**Lemma 9.** *Let T* = *T*_1_ *\* · · · * T*_*m*_*∈𝒯*_*n*_, *where each T*_*i*_*has n*_*i*_*leaves. Then*

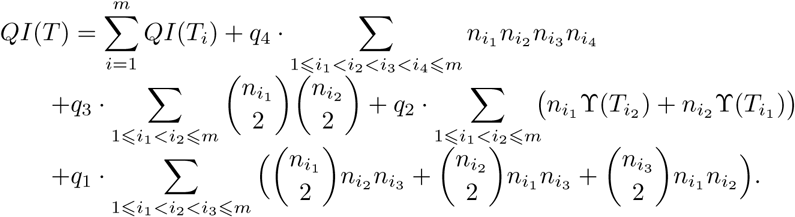

*Proof.* For every *Q ∈ 𝒫*_4_([*n*]), there are the following possibilities:

1. If *Q ⊆L*(*T*_*i*_), for some *i*, then *T* (*Q*) = *T*_*i*_(*Q*). Therefore, the quartets with *Q ⊆L*(*T*_*i*_) contribute *QI*(*T*_*i*_) to *QI*(*T)*.
2. If 3 leaves, say *a, b, c*, in *Q* belong to a subtree *T*_*i*_and the fourth to another subtree *T*_*j*_, then *T* (*Q*):
  - Has shape 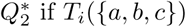 has shape *S*_3_. For every pair of subtrees *T*_*i*_, *T*_*j*_, there are *n*_*j*_r(*T*_*i*_)+ *n*_*i*_r(*T*_*j*_) quartets of this type, and each one of them contributes *q*_2_ to *QI*(*T)*
  - Has shape 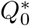 if *T*_*i*_({*a, b, c})* has shape *K*_3_. These quartets do not contribute anything to *QI*(*T)*.
3. If 2 leaves in *Q* belong to a subtree *T*_*i*_and the other 2 to another subtree *T*_*j*_, then *T* (*Q*) has shape 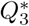. For every pair of subtrees *T*_*i*_, *T*_*j*_, there are 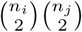 quartets of this type, and each one of them contributes *q*_3_ to *QI*(*T)*.
4. If 2 leaves in *Q* belong to a subtree *T*_*i*_, a third leaf to another subtree *T*_*j*_and the fourth to a third subtree *T*_*k*_, then *T* (*Q*) has shape 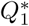. For every triple of subtrees *T*_*i*_, *T*_*j*_, *T*_*k*_, there are 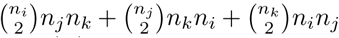quartets of this type, and each one of them contributes *q*_1_ to *QI*(*T)*
5. If each leaf in *Q* belongs to a different subtree *T*_*i*_, then *T* (*Q*) has shape 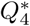. For every quartet of subtrees *T*_*i*_, *T*_*j*_, *T*_*k*_, *T*_*l*_, there are *n*_*i*_*n*_*j*_*n*_*k*_*n*_*l*_ such quartets *Q*, and each one of them contributes *q*_4_ to *QI*(*T)*.

Adding up all these contributions, we obtain the formula in the statement.

**Proposition 10.** *If T ∈ 𝒯*_n_, *QI*(*T) can be computed in time O*(*n*).

*Proof.* Let *T∈𝒯*_n_. Recall that if a certain mapping φ : *V* (*T)→ℝ* can be computed in constant time at each leaf of *T* and in *O*(deg(*v*)) time at each internal node *v* from its value at the children of *v*, then the whole vector (*φ*(*v*))_*v∈V(T)*_), and hence also its sum 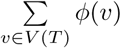 can be computed in *O*(*n*) time by traversing *T* in post-order. Indeed, if we denote by *m*_*k*_ the number of internal nodes of *T* with out-degree *k*, then the cost of computing (*φ*(*v*))_*v∈V(T)*_) through a post-order traversal of *T* is *O (n* +Σ*k m*_*k*_ *· k),* and Σ*k m*_*k*_ *· k* is the number of arcs in *T,* which is at most 2*n -* 2. We shall use t his re mark several times in this proof, and to begin with, we refer to it to recall that the vector *κ*(*v*) _*v∈V* (*T)*_ can be computed in *O*(*n*) time.

Now, in order to simplify the notations, let, for every *v ∈ V*_*int*_(*T)*:

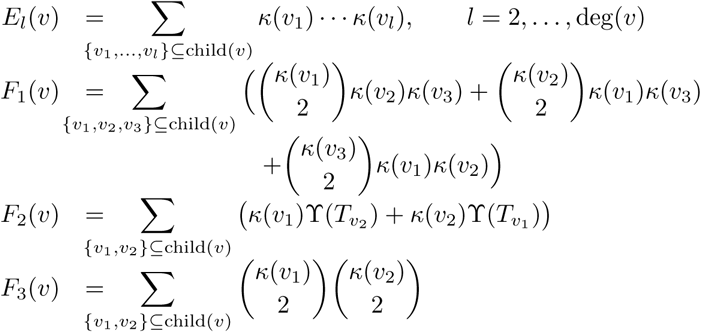

so that

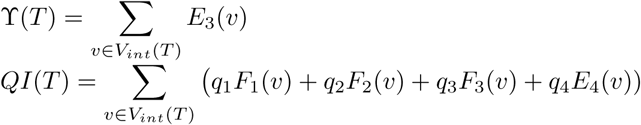

We want to prove now that each one of the vectors

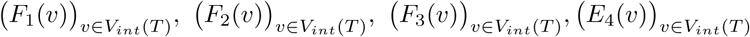

can be computed in *O*(*n*) time, which will clearly entail that *QI*(*T)* can be computed in *O*(*n*) time.

One of the key ingredients in the proof are the *Newton-Girard formulas* [14, *§I.2]:* given a (multi)set of numbers *X* = *{x*_1_,*…, x*_*k*_*}*, if we let, for every *l≥* 1,

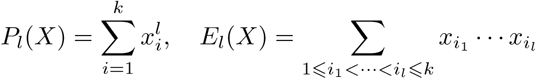

then

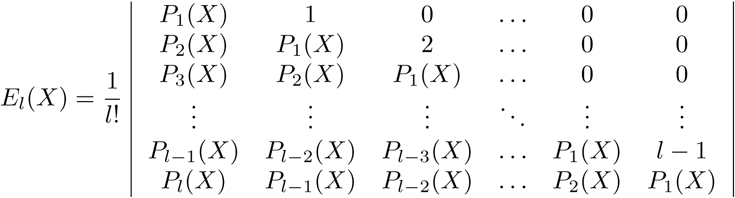

If we consider *l* as a fixed parameter, every *P*_*l*_(*X*) can be computed in time *O*(*k*) and then this expression for *E*_*l*_(*X*) as an *l × l* determinant allows us also to compute it also in time *O*(*k*).

In particular, if, for every *v ∈ V*_*int*_(*V)*, we consider the multiset *X*_*v*_ = *{κ*(*u*) *| u ∈* child(*v*) *}*, then every *E*_*l*_(*v*) = *E*_*l*_(*X*_*v*_) can be computed in time *O*(deg(*v*)) and hence the whole vector *(E*_*l*_*(v))*_*vεVint(T)*_ can be computed in time *O*(*n*). In particular *(E*_*3*_*(v))*_*vεVint(T)*_ and *(E*_*4*_*(v))*_*vεVint(T)*_ can be computed in linear time.

Then, using the recursion

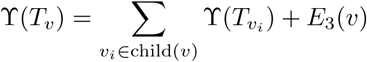

we deduce that the whole vector *(r(T*_*v*_))_*vεVint(T)*_ can also be computed in time *O*(*n*). And then, since

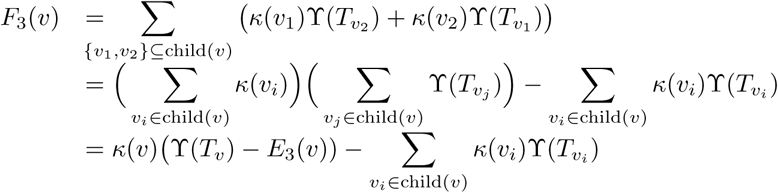

this implies that each *F*_3_(*v*) can be computed in time *O*(deg(*v*)) and hence that the whole vector *(F*_*3*_*(v))*_*vεVint(T)*_ can be computed in time *O*(*n*).

Let us focus now on

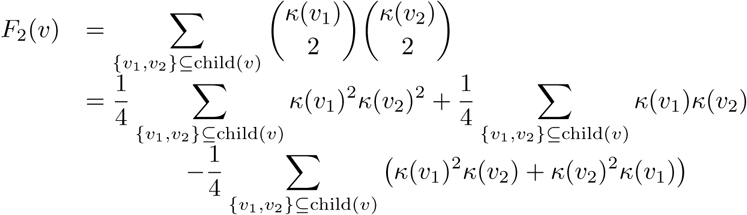

In this expression,

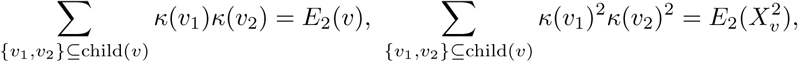

Where 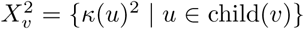, and hence they are computed in time *O*(deg(*v*)). As far as the subtrahend goes,

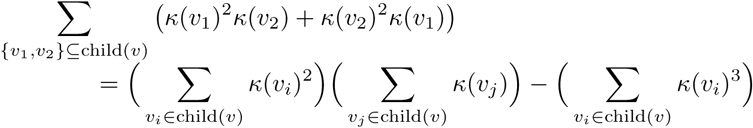

and hence it can also be computed in time *O*(deg(*v*)). Therefore, the whole vector *(F*_*2*_*(v))*_*vεVint(T)*_ can be computed in time *O*(*n*).

Let us consider finally

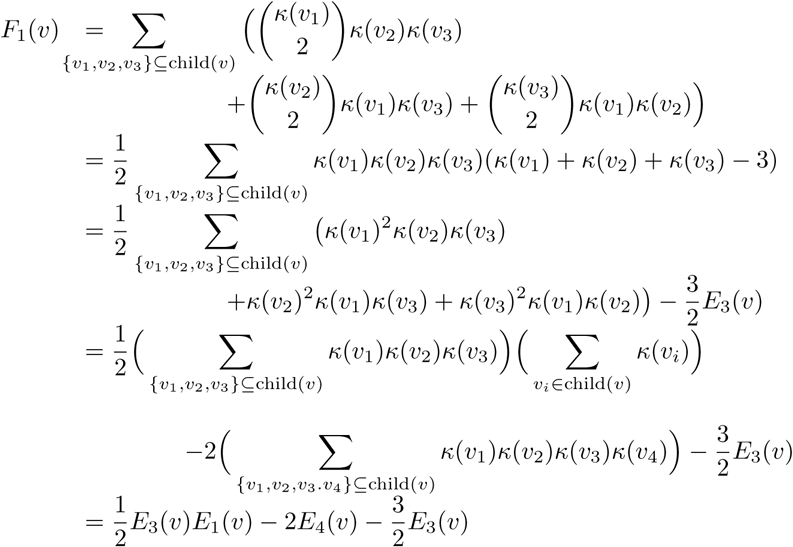

Hence, it can also be computed in time *O*(deg(*v*)) and therefore, the whole vector *(F*_*1*_*(v))*_*vεVint(T)*_ can be computed in time *O*(*n*).

## 4. Trees with maximum and minimum *QI*

Let *n* ≥ 4. In this section we determine which trees in 𝒯_*n*_ and ℬ𝒯_*n*_ have the largest and smallest corresponding quartet indices. The multifurcating case is easy:

**Theorem 11.** *The minimum value of QI in𝒯*_*n*_ *is reached exactly at the combs K*_*n*_, *a nd it is 0. The maximum value value of QI in 𝒯*_*n*_ *is reached exactly at the star S*_*n*_, *and it is* 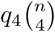

*Proof.* Since the *QI*-value of a quartet goes from 0 to *q*_4_, we have that 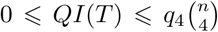 for every *T ∈𝒯* _*n*_. Now, all quartets displayed by a comb *K*_n_ have shape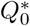, and therefore *QI*(*K*_n_) = 0, while all quartets displayed by a star *S*_n_ have shape 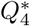, and therefore 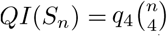

As far as the uniqueness of the trees yielding the maximum and minimum values of *QI* goes, notice that, on the one hand, if *T* is not a comb, then it displays some quartet of shape other than 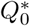, because it contains either some multifurcating internal node, which becomes the root of some multifurcating quartet, or two cherries that determine a quartet of shape 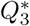. This implies that if *T* ≠ *K*_*n*_, then *QI*(*T) >* 0. On the other hand, if *T ‡S*_n_, then its root has some child that is not a leaf and therefore *T* displays some quartet of shape other than 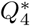, which implies that 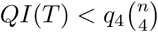.

Therefore, the range of *QI* on *𝒯*_*n*_ goes from 0 to 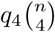. This is one order of magnitude wider than the range of the total cophenetic index, which, going from 0 to 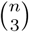 was so far the balance index in the literature with the widest range [17].

It remains to characterize those *bifurcating* phylogenetic trees with largest *QI*, or, equivalently, with largest *QIB*. They turn out to be exactly the maximally balanced trees, as defined at the end of §2.1. The proof is similar to that of the characterization of the bifurcating phylogenetic trees with minimum total cophenetic index provided in [17, §4].

**Lemma 12.** *Let T ∈ 𝔐 𝒯*_*n*_ *be the bifurcating phylogenetic tree depicted in Fig 5.(a). For every i* = 1, 2, 3, 4, *let n*_*i*_ = *|L*(*T*_*i*_)*|, and assume that n*_1_ *> n*_3_ *and n*_2_ *> n*_4_. *Then, QIB*(*T) is not maximum in 𝔐 𝒯*_*n*_.

**Figure 5:**
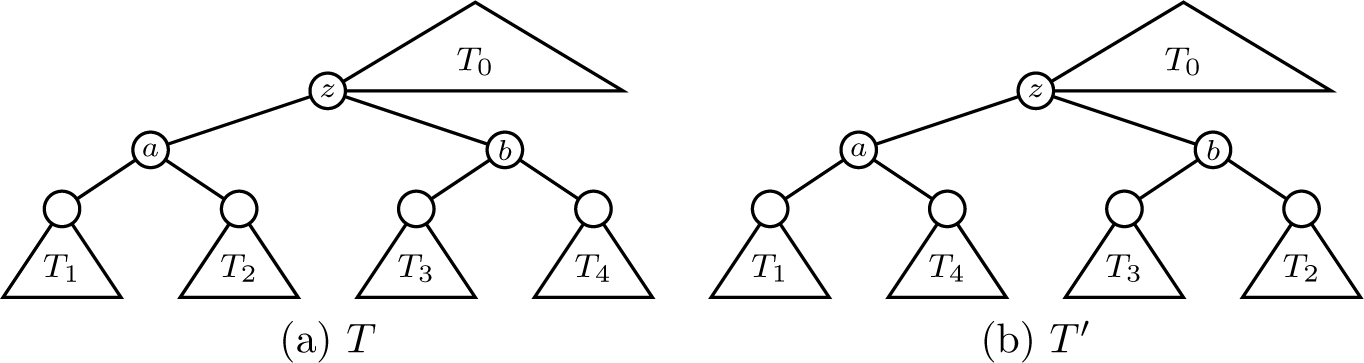
(a) The tree *T* in the statement of Lemma 12. (b) The tree *T* ^*′*^in the proof of Lemma 12.

*Proof.* Let *T* ^*′*^ be the tree obtained from *T* by interchanging *T*_2_ and *T*_4_; see Fig 5.(b). We shall prove that *QIB*(*T* ^*′*^) *> QIB*(*T)*.

Let Σ_z_ be the set of labels of *T*_z_, which is also the set of labels of 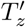 To simplify the language, we shall understand the common subtree *T*_0_ of *T* and *T* ^*′*^ as a phylogenetic tree on ([*n*] *\ Σ*_z_) *∪* {*z*}. Then, for every *Q* = {*a, b, c, d*} *∈ 𝒫*_4_([*n*]):

- If *Q ∩Σ*_z_ – *Ø*, then *T* (*Q*) – *T* ^*′*^(*Q*) – *T*_0_(*Q*).
- If *Q ∩* Σ_z_ is a single label, say *d*, then *T* (*Q*) – *T* ^*′*^(*Q*) – *T*_0_(*{a, b, c, z}*).
- If *Q∩*Σ_z_ consist of two labels, say *c, d*, then *T* (*Q*) – *T* ^*′*^(*Q*). More specifically: *T* (*Q*) –*T* ^*′*^(*Q*) – ((*a, b*), (*c, d*)) when *T*_0_({*a, b, z})* = ((*a, b*), *z*); *T* (*Q*) – *T* ^*′*^(*Q*) – (*a,* (*b,* (*c, d*))) when *T*_0_({ *a, b, z})* = (*a,* (*b, z*)); and *T* (*Q*) – *T* ^*′*^(*Q*) – (*b,* (*a,* (*c, d*))) when *T*_0_({*a, b, z})* = (*b,* (*a, z*)).
- If *Q ∩* Σ_*z*_ consist of three labels, then *T* (*Q*) and *T* ^*′*^(*Q*) are both combs.

Therefore, *T* (*Q*) and *T* ^*′*^(*Q*) can only be different when *Q ⊆* Σ_*z*_, in which case *T* (*Q*) = *T*_*z*_(*Q*) and 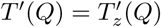. This implies that

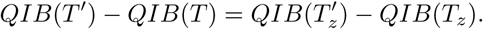

Now, to compute the difference in the right hand side of this equality, we apply Lemma 7:

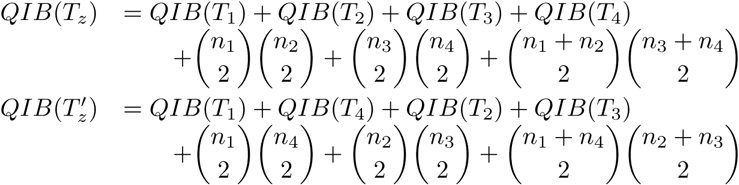

and hence

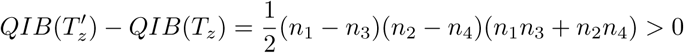

because *n*_1_ *> n*_3_ and *n*_2_ *> n*_4_ by assumption.

**Lemma 13.** *Let T ∈ 𝔅 𝒯* _*n*_ *be a bifurcating phylogenetic tree containing a leaf whose sibling has at least 3 descendant leaves. Then, QIB*(*T) is not maximum in 𝔅 𝒯* _*n*_.

**Figure 6:**
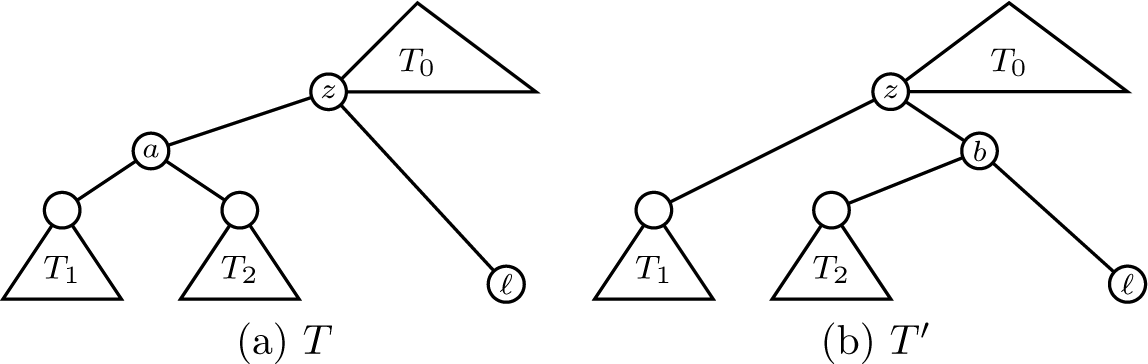
(a) The tree *T* in the statement of Lemma 13. (b) The tree *T* ^*′*^ in the proof of Lemma 13.

*Proof.* Assume that the tree *T ∈ 𝔅 𝒯* _*n*_ in the statement is the one depicted in Fig 6.(a), with *l* a leaf such that the subtree *T*_*a*_ rooted at its sibling *a* has |*L*(*T*_*a*_) | ⩾ 3. Let *n*_1_ = |*L*(*T*_1_) | and *n*_2_ = |*L*(*T*_2_)| and assume *n*_1_ ⩾ *n*_2_: then, since *n*_1_ + *n*_2_ ⩾3, *n*_1_ ⩾2. Let then *T* ^*′*^ be the tree depicted in Fig 6.(b): we shall prove that *QIB*(*T* ^*′*^) *> QIB*(*T)*. Reasoning as in the proof of the last lemma, we deduce that

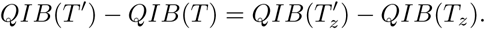

Now, using Lemma 7, we have that

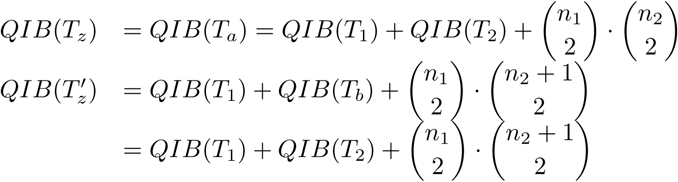

and therefore

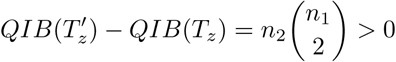

as we wanted to prove.

**Theorem 14.** *For every T∈ 𝔅 𝒯*_*n*_, *QIB*(*T) is maximum in 𝔅 𝒯*_*n*_ *if, and only if, T is maximally balanced.*

*Proof.* Assume that *QIB*(*T)* is maximum in *𝔅 𝒯*_*n*_ but that *𝒯∈ 𝔅 𝒯*_*n*_ is not maximally balanced, and let us reach a contradiction. Let *z* be a non-balanced internal node in *T* such that all its proper descendant internal nodes are balanced, and let *a* and *b* be its children, with *κ*(*a*) ⩾ *κ*(*b*) + 2.

If *b* is a leaf, then, by Lemma 13, *QIB*(*T)* cannot be maximum in *𝔅 𝒯*_*n*_. Therefore, *a* and *b* are internal, and hence balanced. Let *v*_1_, *v*_2_ be the children of *a*, *v*_3_, *v*_4_ the children of *b*, and *n*_*i*_ = *κ*(*v*_*i*_), for *i* = 1, 2, 3, 4. Without any loss of generality, we shall assume that *n*_1_ ⩾ *n*_2_ and *n*_3_ ⩾*n*_4_. Then, since *a* and *b* are balanced, *n*_1_ = *n*_2_ or *n*_2_ + 1 and *n*_3_ = *n*_4_ or *n*_4_ + 1. Then,*n*_1_ + *n*_2_ – *κ*(*a*) ⩾*κ*(*b*) + 2 = *n*_3_ + *n*_4_ + 2 implies that *n*_1_ *> n*_3_.

Now, by Lemma 12, since by assumption *QIB*(*T)* is maximum on *𝔅 𝒯* _*n*_, it must happen that *n*_1_ *> n*_3_ ⩾*n*_4_ ⩾ *n*_2_. This forbids the equality *n*_1_ = *n*_2_, and hence *n*_1_ −1 – *n*_2_ – *n*_3_ – *n*_4_. But this contradicts that *n*_1_ + *n*_2_ ⩾*n*_3_ + *n*_4_ + 2.

This implies that a non maximally balanced tree in *𝔅 𝒯*_*n*_ cannot have maximum *QIB*, and therefore the maximum *QIB* in *𝔅 𝒯*_*n*_ is reached at the maximally balanced trees, which have all the same shape and hence the same *QIB* index.

So, the only bifurcating trees with maximum *QIB* (and hence with maximum *QI*) are the maximally balanced. This maximum value of *QIB* on *𝔅 𝒯*_*n*_ is given by the following recurrence.

**Lemma 15.** *For every n, let q*_n_ *be the maximum of QIB on 𝔅 𝒯*_*n*_. *Then, q*_1_ – *q*_2_ – *q*_3_ – 0 *and*

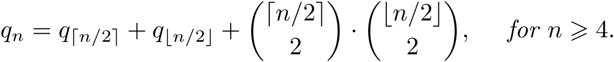

*Proof.* This recurrence for *q*_*n*_ is a direct consequence of Lemma 7 and the fact that the root of a maximally balanced tree in *𝔅 𝒯*_*n*_ is balanced and the subtrees rooted at their children are maximally balanced.

The sequence (*q*_*n*_)_*n*_ seems to be new, in the sense that it has no relation with any sequence previously contained in Sloane’s *On-Line Encyclopedia of Integer Sequences* [25]. *Its values for n* – 4,*…,* 20 are

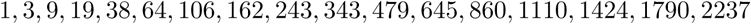

It is easy to prove, using the Master theorem for solving recurrences [8, Thm. 4.1], that *q*_*n*_ grows asymptotically in *O*(*n*^*4*^). Moreover, it is easy to compute *q*_2_*n* from this recurrence, yielding

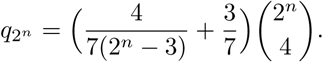

In particular, 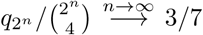, which is in agreement with the probability of ((*\*, \**), (*\*, \**)) under the *β*-model when β *→ ∞*; cf. [1, §4.1].

## 5. The expected value and the variance of *QI*

Let *𝔐* – (*P*_*n*_)_*n*_ be a probabilistic model of phylogenetic trees and *QI*_*n*_ the random variable that chooses a phylogenetic tree *T* ∈ *𝒯*_n_ with probability distribution *P*_*n*_ and computes *QI*(*T)*. In this section we are interested in obtaining expressions for the expected value *E*_*𝔐*_ (*QI*_*n*_) and the variance *V ar*_*𝔐*_ (*QI*_*n*_) of *QI*_*n*_ under suitable models. *𝔐.*

Next lemma shows that, to compute these values, we can restrict ourselves to work with unlabeled trees. Let 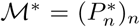 the probabilistic model of trees induced by *𝔐* and 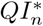 the random variable that chooses a tree 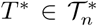 with probability distribution 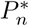 and computes *QI*(*T* ^*^), defining it as *QI*(*T)* for some phylogenetic tree *T* of shape *T* ^*^.

**Lemma 16.** *For every n ⩾*1, *the distributions of QI*_*n*_ *and* 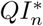 *are the same. In particular, their expected values and their variances are the same.*

*Proof.* Let *f*_*q*_*I*_*n*_ and 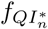 be the probability density functions of the discrete random variables *QI*_*n*_ and 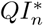, respectively. Then, for every *x*_0_ ∈ ℝ,

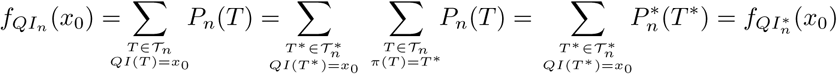

**Proposition 17.** *If 𝔐* * *is sampling consistent, then*

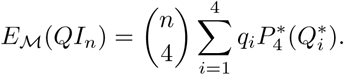

*Proof.* By Lemma 16, *E*_𝔐_ (*QI*_*n*_) is equal to the expected value 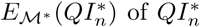 under *𝔐* *, which can be computed as follows:

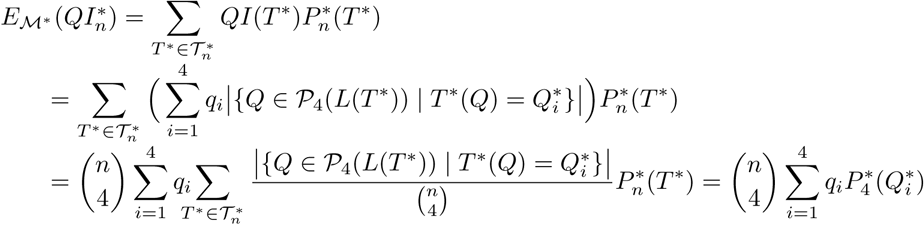

because, for every *i* – 1,*…,* 4,

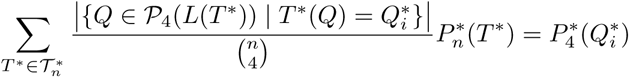

by the sampling consistency of *𝔐* *.

The α-γ model for unlabeled trees 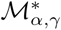 is sampling consistent [6, Prop. 12]. Therefore, applying the last proposition using the distribution 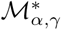 on 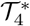 given in §2.2, we have the following result.

**Corollary 18.** *Let 𝔐*_α,γ_ = (*P*_*α,n*_)_*n ⩾* 1_ *be the α-γ-model on* (*𝒯*_*n*_)_*n ⩾* 1_, *with α ∈* [0, 1]. *Then*

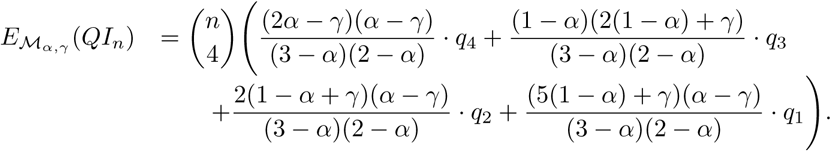

If *𝔐* – (*P*_n_)_n_ is a probabilistic model of bifurcating phylogenetic trees, so that 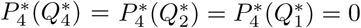, then the expression in Prop. 17 becomes

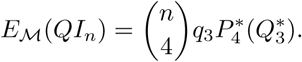

Making *q*_3_ – 1, we obtain the following result.

**Corollary 19.** *If 𝔐 is a probabilistic model of bifurcating phylogenetic trees such that 𝔐*\* *is sampling consistent, then*

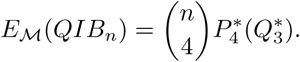

Since the *α* and β-models of bifurcating (unlabeled) trees are sampling consistent, this corollary together with the probabilities of 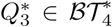 under these models given in §2.2 entail the following result.

**Corollary 20.** *Let* 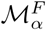 *be Ford’s α-model for bifurcating phylogenetic trees, with α ∈* [0, 1], *and let 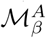 be Aldous’ β-model for bifurcating phylogenetic trees, with β* ∈] *-* 2, *∞* [. *Then:*

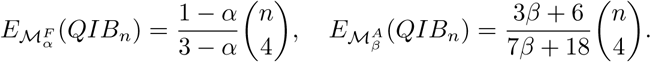

It is easy to check that 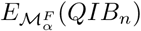 agrees with 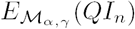 (up to the factor *q*_3_) when α – γ.

In particular, under the Yule model, which corresponds to α – 0 or β – 0, and the uniform model, which corresponds to α – 1*/*2 or β = *-*3*/*2, the expected values of *QIB*_*n*_ are, respectively,

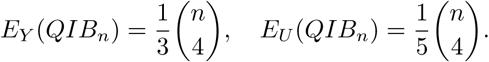

Let us deal now with the variance of *QI*_*n*_. To simplify the notations, for every *k* = 5, 6, 7, 8, for every 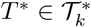 and for every *i, j* ∈ {1, 2, 3, 4}, let

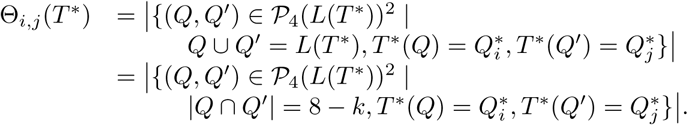

Notice that Θ_i,j_(*T* ^*^) – Θ_*j,i*_(*T* ^*^).

**Proposition 21.** *If 𝔐*\* *is sampling consistent, then*

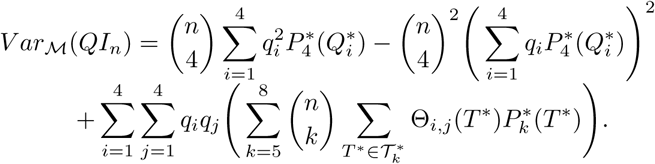

*Proof.* Since, by Lemma 16, 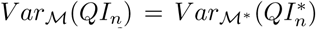 we shall compute the latter using the formula 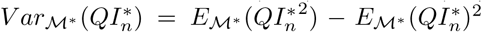 and therefore we need to compute 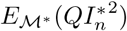

For every 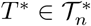, for every 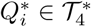 and for every *Q ∈ 𝒫*_4_(*L*(*T* ^*^)), set

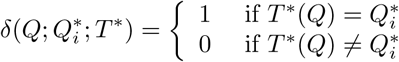

Then:

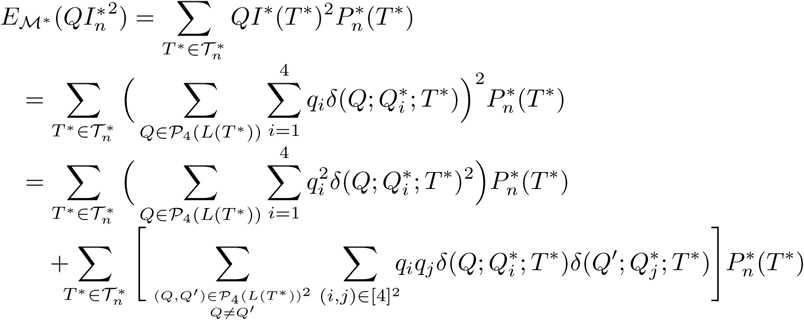

Now, since 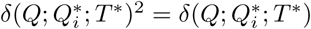,

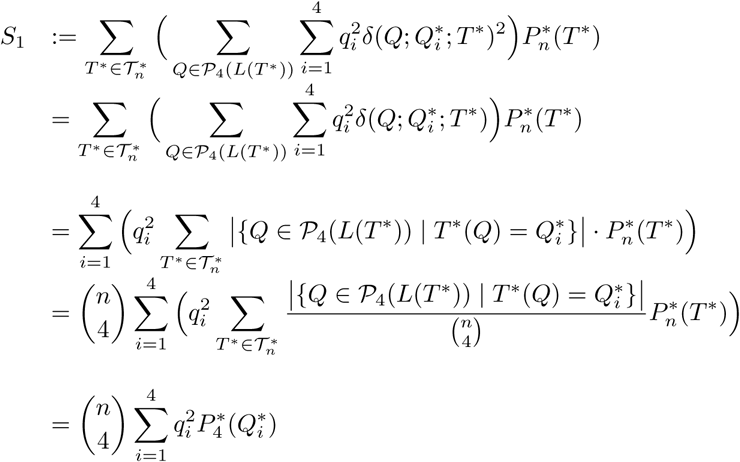

by the sampling consistency of *𝔐*\*.

As far as the second addend in the previous expression for 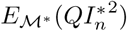 goes, we have

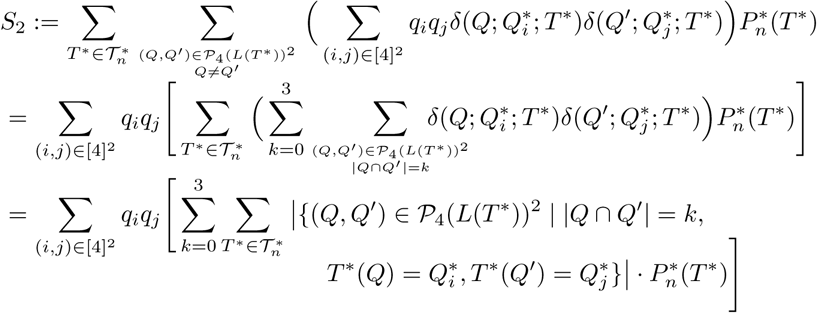

Now notice that, for every *k* – 0,*…,* 3,

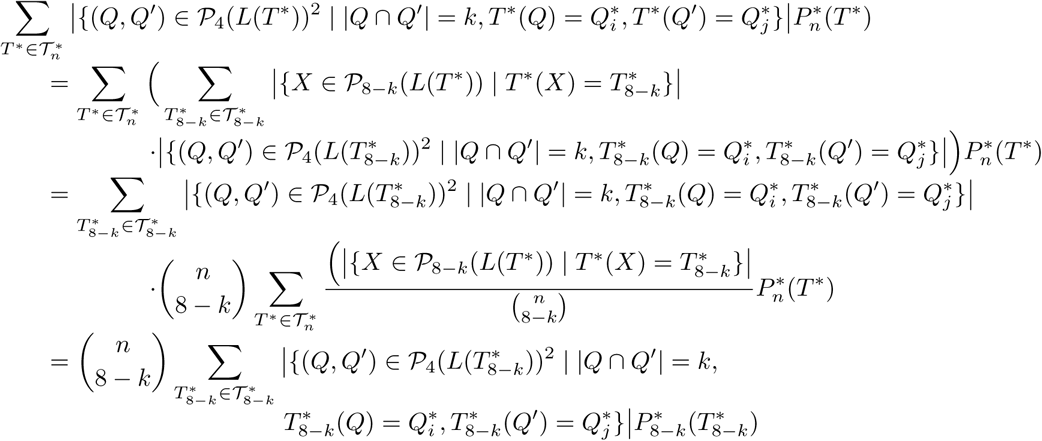

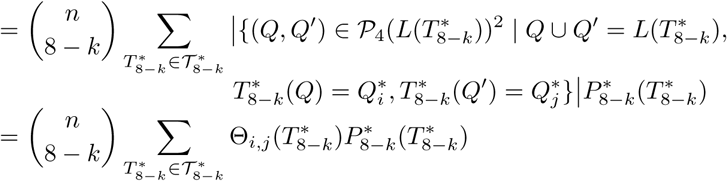

again by the sampling consistency of *𝔐*\*. Therefore,

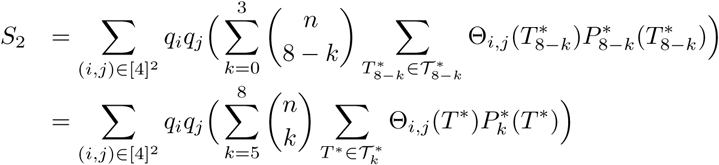

The formula in the statement is then obtained by writing 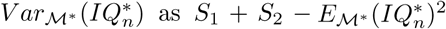 and using the expression for 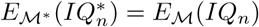 given in Proposition 17.

Again, if *𝔐* = (*P*_n_)_n_ is a probabilistic model of bifurcating phylogenetic trees, so that 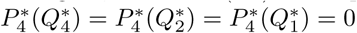 then,taking *q*_3_ = 1,this proposition impiles that

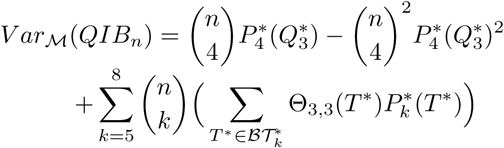

In this bifurcating case, the figures Θ_3,3_(*T* ^*^) appearing in this expression can be easily computed by hand: they are provided in Table 2 in the Appendix A.4. We obtain then the following result.

**Corollary 22.** *If 𝔐is a probabilistic model of bifurcating phylogenetic trees such that 𝔐* * *is sampling consistent, then, with the notations for trees given in Table 2 in the Appendix A.4,*

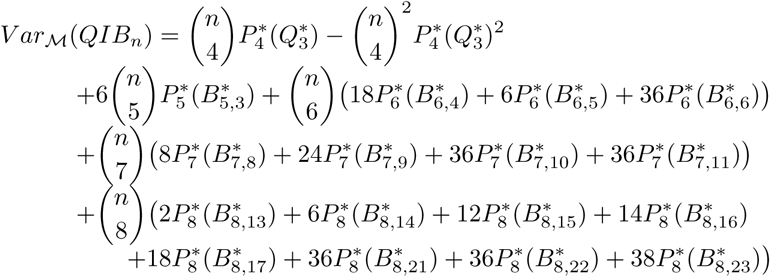

Proposition 21 and Corollary 22 reduce the computation of *V ar*_*𝔐*_(*QI*_*n*_) or *V ar*_*𝔐*_(*QIB*_*n*_) to the explicit knowledge of 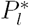 for *l* – 4, 5, 6, 7, 8. In particular, they allow to obtain explicit formulas for the variance of *QIB*_n_ under the α and the β-models, and for the variance of *QI*_n_ under the α-γ-model.

As far as the bifurcating case goes, on the one hand, the probabilities under the *α*-model of the trees appearing explicitly in the formula for the variance of *QIB*_*n*_ in Corollary 22 are those given in Table 3 in the Appendix A.4 (they are explicitly computed in [15, Suppl. Mat.]). Plugging them in the formula given in Corollary 22 above, we obtain the following result.

**Corollary 23.** *Under the α-model,*

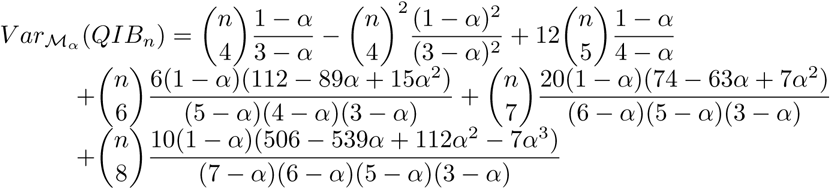

In particular, the leading term in *n* of 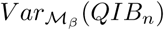 is

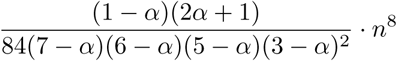

On the other hand, the probabilities under the β-model of the same trees are given in Table 4 in the Appendix A.4, yielding the following result.

**Corollary 24.** *Under the β-model,*

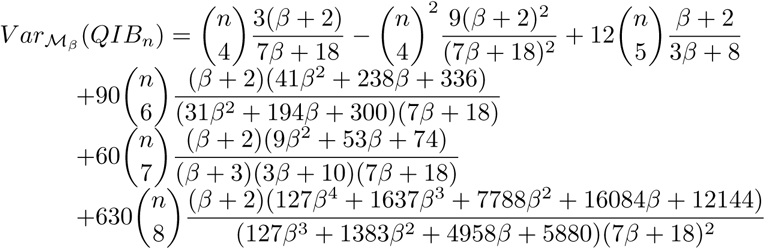

In particular, the leading term in *n* of 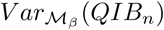 is

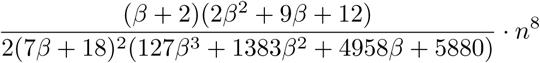

When α – 0 or β – 0, which correspond to the Yule model, both formulas for the variance of *QIB*_*n*_ reduce to

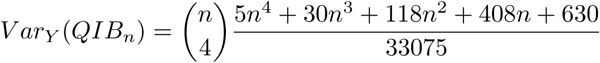

In the Appendix A.4 we give an independent derivation of this formula, which provides extra evidence of the correctness of all these computations.

As far as the uniform model goes, when *α* = 1*/*2 or β = 0, these formulas yield

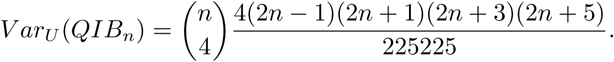

Finally, as far as the *α*-*γ*-model goes, we have written a set of Python scripts that compute all Θ_i,j_(*T* ^*^), *i, j* = 1, 2, 3, 4, as well as 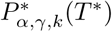 for every 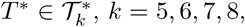, *k* – 5, 6, 7, 8, and combine all these data into an explicit formula for 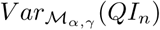. The Python scripts and the resulting formula (in text format and as a Python script that can be applied to any values of *n*, α, and ϒ) can be found in the GitHub page https://github.com/biocom-uib/Quartet_Index companion to this paper. In particular, the plain text formula (which is too long and uninformative to be reproduced here) is given in the document variance_table.txt therein. It can be easily checked using a symbolic computation program that when α – γ it agrees with the variance under the α-model given in Corollary 23.

## 6. Conclusions

In this paper we have introduced a new balance index for phylogenetic trees, the quartet index *QI*. This index makes sense for polytomic trees, it can be computed in time linear in the number of leaves, and it has a larger range of values than any other shape index defined so far. We have computed its maximum and minimum values for bifurcating and arbitrary trees, and we have shown how to compute its expected value and variance under any probabilistic model of phylogenetic trees that is sampling consistent and invariant under relabelings. This includes the popular uniform, Yule, α, β and α-ϒ-models. This paper is accompanied by the GitHub page https://github.com/biocom-uib/Quartet_Index where the interested reader can find a set of Python scripts that perform several computations related to this index.

We want to call the reader’s attention on a further property of the quartet index: it can be used in a sensible way to measure the balance of *taxonomic trees*, defined as those rooted trees of fixed depth (but with possibly out-degree 1 internal nodes) with their leaves bijectively labeled in a set of taxa. The usual taxonomies with fixed ranks are the paradigm of such taxonomic trees. It turns out that the classical balance indices cannot be used in a sound way to quantify the balance of such trees. For instance, Colless’ index cannot be applied to non-bifurcating trees, and Sackin’s index, being the sum of the depths of the leaves in the tree, is constant on all taxonomic trees of fixed depth and number of leaves. As far as the total cophenetic index, it is straightforward to check from its very definition that the taxonomic trees with maximum and minimum total cophenetic values among all taxonomic trees of a given depth and a given number of leaves are those depicted in Fig 7, which, in our opinion, should be considered as equally balanced. We believe that the *QI* index can be used to capture the symmetry of a taxonomic tree in a natural way, and we hope to report on it elsewhere.

**Figure 7:**
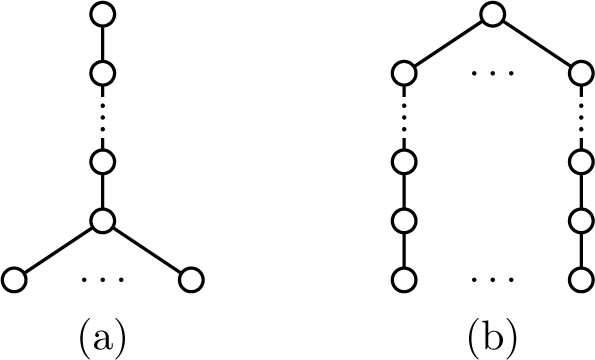
The shapes of the taxonomic trees with maximum (a) and minimum (b) total cophenetic values among all taxonomic trees of given depth and number of leaves.

## Acknowledgements

A preliminary version of this paper was presented at the *Workshop on Algebraic and combinatorial phylogenetics* held in Barcelona (June 26–30, 2017). We thank Mike Steel and Seth Sullivant for their helpful suggestions on several aspects of this paper. This research was partially supported by the Spanish Ministry of Economy and Competitiveness and the European Regional Development Fund through project DPI2015-67082-P (MINECO/FEDER).

## Appendix: Some computations

### *A.1: Computation of* 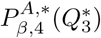

We recall from [1] the definition of the probability *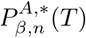* of a tree 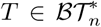 under the *β*-model. Let β *∈*] *-* 2, *∞* [. For every *m ⩾* 2 and *a* – 1,*…, m -* 1, let

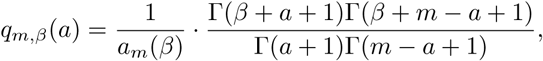

where *a*_*m*_(β) is a suitable normalizing constant so that 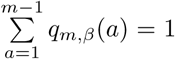. For every *m ⩾* 2 and *a* – 1,*…, lm/*2*J*, let

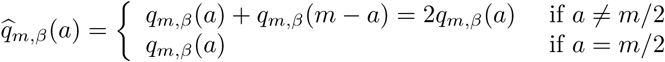

Then, for a given desired number *n ⩾* 1 of leaves:

1. Start with a tree 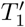 consisting of a single node labeled *n*. Let 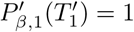
2. At each step *j* – 1,*…, n –* 1, the current tree 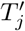 contains leaves with label greater than 1. Then, choose equiprobably a leaf in 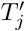 with a label *m* greater than 1, choose a number *a* – 1,*…, lm/*2*J* with probability distribution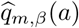, and split this leaf into a cherry with a child labeled *a* and a child labeled *m - a*. The resulting tree 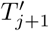 has then probability

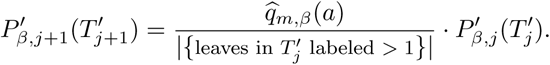
3. When the desired number *n* of leaves is reached, all leaves are labeled 1 and 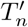 can be understood as a tree. Then, the probability of a given tree is defined as the sum of the probabilities of all ways of obtaining it by means of the previous procedure; that is, for every 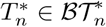,its probability under the β-model is

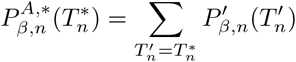

**Lemma 25.** *For every β ∈*] *-* 2, *∞* [,

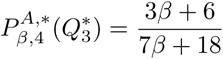

*Proof.* To compute 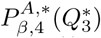 we start with a single node labeled 4. In order to obtain a maximally balanced tree ((1, 1), (1, 1)) using the previous procedure, in the first step we must split this node into a cherry with both leaves labeled 2. The probability of choosing this split is

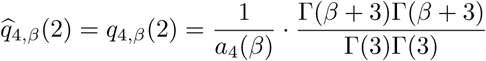

Let’s compute the normalizing constant *a*_4_(β): since

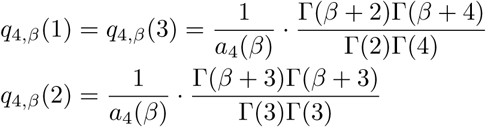

imposing that *q*_4,*β*_(1) + *q*_4,*β*_(2) + *q*_4,*β*_(3) – 1 we obtain

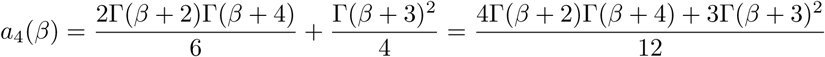

Therefore,

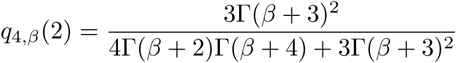

In the second step, we choose one of the leaves with probability 1*/*2 and we split it into a cherry (1, 1). Since there is only one way of splitting a leaf labeled 2, *q*_2,*β*_(1) = 1. So, the probability of the tree obtained in this step is

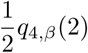

Then, in the third step, we are forced to choose the other leaf labeled 2 and to split it into a cherry (1, 1). We obtain a maximally balanced tree with all its leaves labeled 1 and its probability is still *q*_4,*β*_(2)*/*2.

Now, there are two ways of obtaining the tree ((1, 1), (1, 1)) with this construction, depending on which leaf of the cherry (2, 2) we choose to split first. So, the probability of the tree 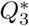 is

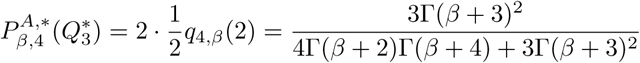

Finally, using that Γ(*x* + 1) = *x* Γ (*x*), we have that

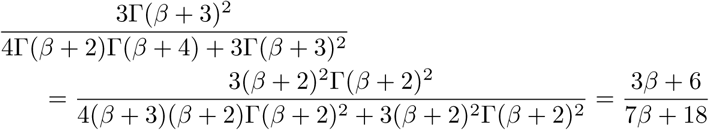

as we claimed.

### A.2: Computation of 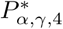

We recall from [6] the definition of the probability 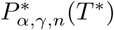 of a tree 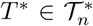 under the *α*-γ-model. Let 0 ⩽ γ *⩽ α* ⩽ 1 and *n ⩾* 1.

1.Start with the tree *T*_1_ ∈ *𝒯* _1_ consisting of a single node labeled 1. Let *P*_α,γ,1_ (*T*_1_) – 1.

*2.*For every *m* – 1,*…, n –* 1, let *T m*+1 *ϵ 𝒯* _*m*+1_ be obtained by adding a new leaf labeled *m* + 1 to *T*_*m*_. Then:

*If *T*_*m*+1_ is obtained from *T*_*m*_ by adding the new leaf to an arc *e*, then:

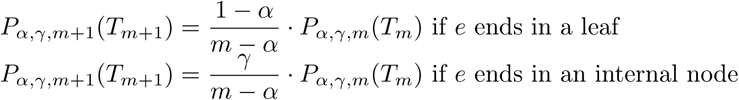

*If *T*_*m* +1_ is obtained from *T*_*m*_ by adding the new leaf to a new root, then

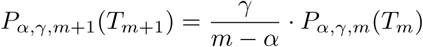

*If *T*_*m* +1_ is obtained from *T*_*m*_ by adding the new leaf as a child of an internal node *u*, then

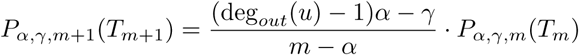

3.When the desired number *n* of leaves is reached, the probability 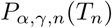 of the resulting tree *T*_*n*_ is the one obtained in this way. Then, the probability 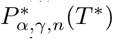 of a given tree *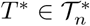* is defined as the sum of the probabilities of all phylogenetic trees with that shape

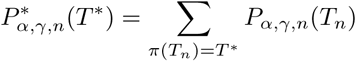

**Lemma 26.** *With the notations of Figure 3:*

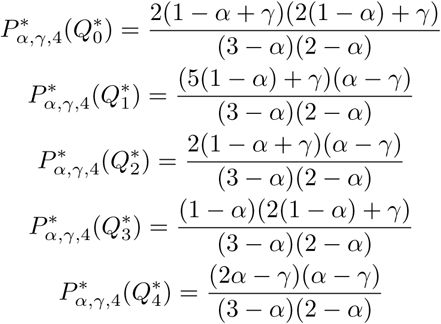

*Proof.* To compute these probabilities, we shall already start with the cherry *T*_2_ = (1, 2) in 𝒯 _2_, which has probability *P*_α,γ,2_ (*T*_2_) = 1. Every tree in 𝒯 _3_ is obtained by adding a leaf labeled 3 to *T*_2_. These trees are described in Figure 8. Their probabilities are:

*S*_3_ is obtained by adding the leaf 3 to the root of *T*_2_. Its probability is then

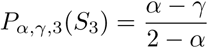

*K*^(^1^)^ and *K*^(^2^)^ are obtained by adding the leaf 3 to an arc in *T*_2_. Their probability is then

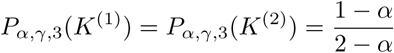

*K*^(^3^)^ is obtained by adding the leaf 3 to a new root. Its probability is then

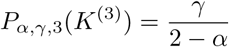

**Figure 8:**
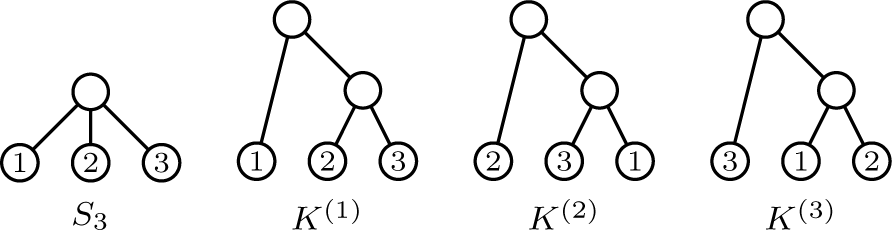
The phylogenetic trees in *𝒯*3.

Let us move finally to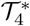:

A tree of shape 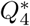 can only be obtained by adding the leaf 4 to the root of the tree *S*_3_. Its probability is, then,

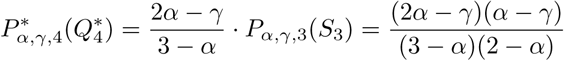

A tree of shape 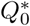 can be obtained by adding the leaf 4 in some tree 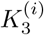 either to a new root, to the arc from the root to the other internal node, or to one of the arcs in its cherry. Its probability is, then,

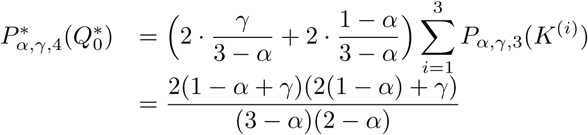

A tree of shape 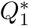 can be obtained by adding the leaf 4 either to one of the three arcs in the tree *S*_3_ or to the root of some tree 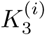. Its probability is, then,

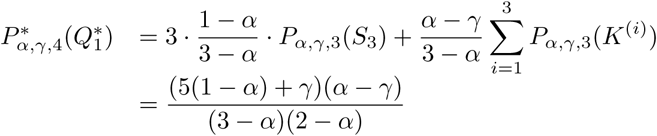

A tree of shape 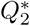 can be obtained by adding the leaf 4 either to a new root in the tree *S*_3_ or to the non-root internal node in some tree 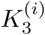. Its probability is, then,

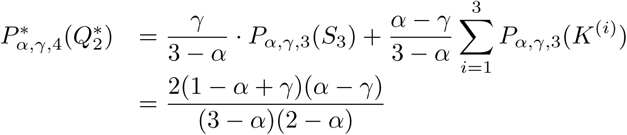

A tree of shape 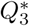 can only be obtained by adding the leaf 4 to the arc from the root to its only leaf child in some tree 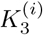. Its probability is, then,

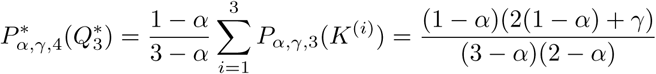

### A.3: An alternative derivation of the variance of QIB_*n*_ under the Yule model

In this section we give an alternative proof of the following result.

**Proposition 27.** *Under the Yule model,*

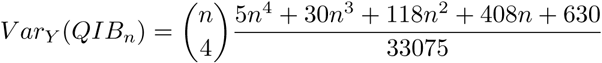

*Proof.* By Lemma 7, *QIB* on 𝔅 𝒯_n_ is a bifurcating recursive tree shape statistic satisfying the recurrence

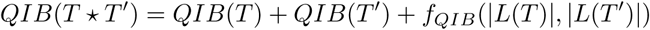

with 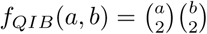. Then, it satisfies the hypothesis in [4, Cor. 1] with

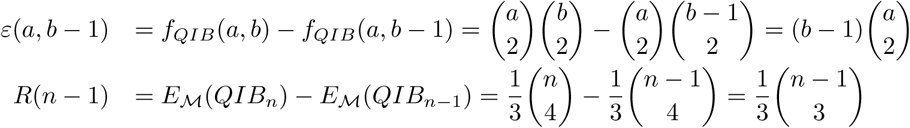

Since *E*_*Y*_ (*QIB*_1_) = 0 and *f*_*QIB*_(*n –* 1, 1) = 0, applying the aforementioned result from [4] we have that

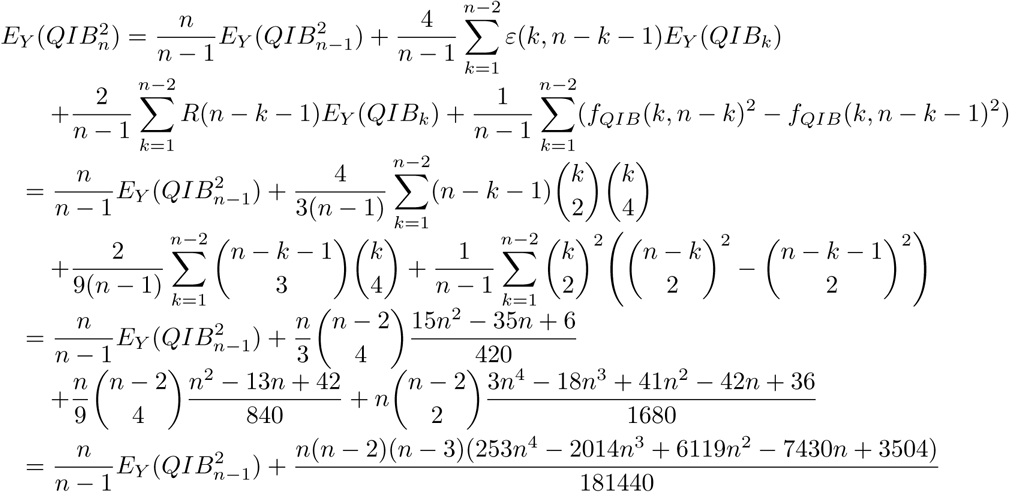

Dividing by *n* both sides of this expression for 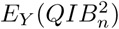 and setting 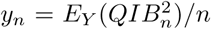, we obtain the recurrence

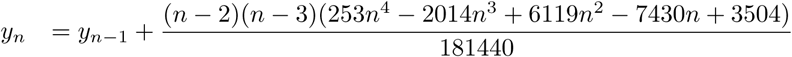

Since *y*_0_ – *y*_1_ – 0, its solution is

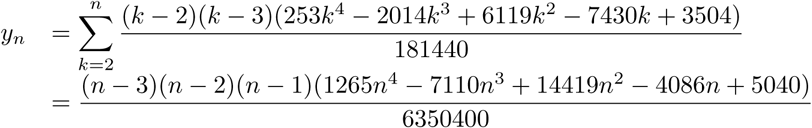

from where we obtain

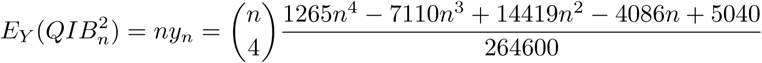

Finally

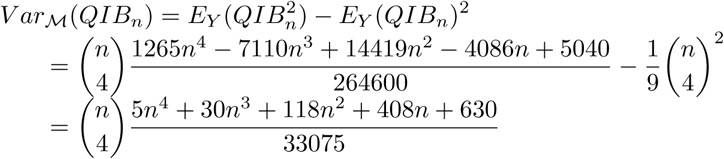

as we claimed.

### A.4: Some tables used in §5

**Table 2:**
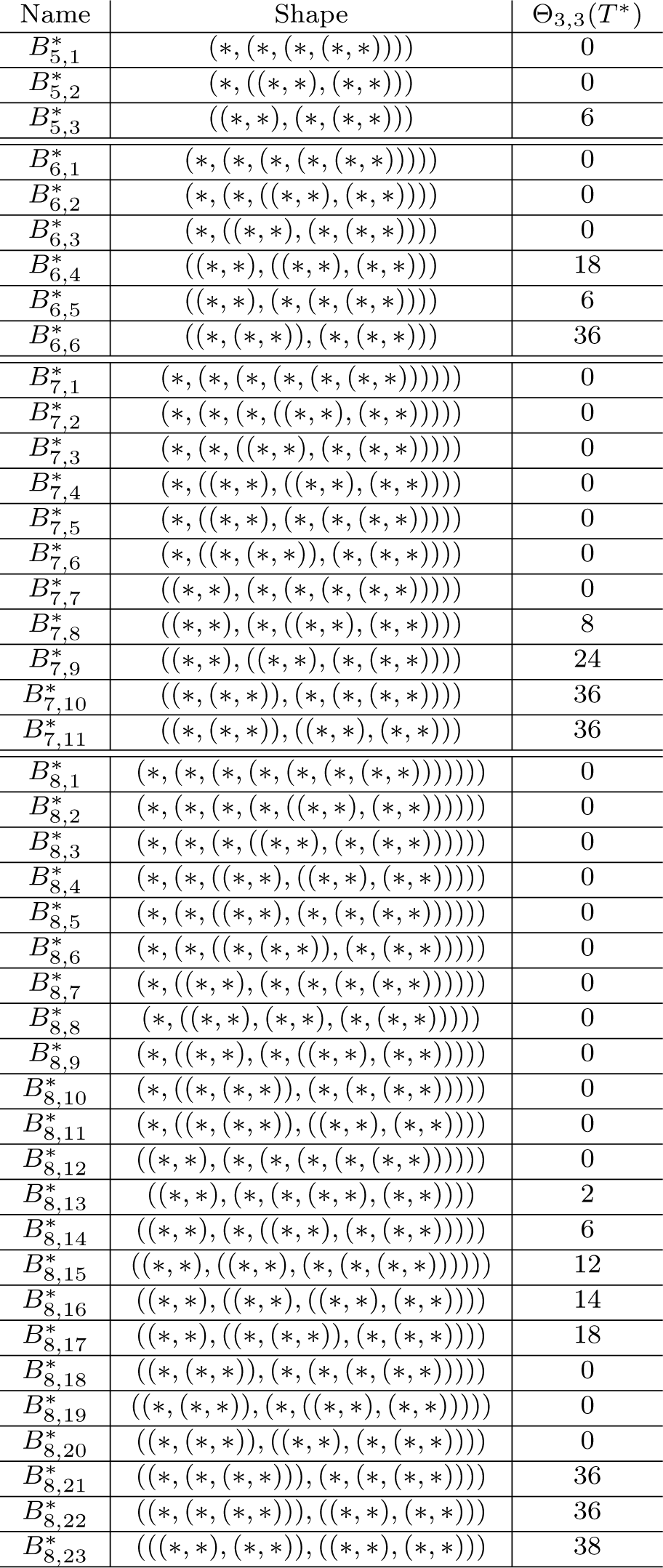
Coefficients of the probabilities of the trees in 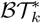, for *k* = 5, 6, 7, 8, in the formula for the variance of *QIB*_n_.

**Table 3:** Probabilities under the α-model of the trees involved in the formula for the variance of *QIB*_n_

| Tree | $P_{\alpha,n}^{A,*}$ |
| --- | --- |
| $Q_3^*$ | $\frac{1-\alpha}{3-\alpha}$ |
| $B_{5,3}^*$ | $\frac{2(1-\alpha)}{4-\alpha}$ |
| $B_{6,4}^*$ | $\frac{(1-\alpha)^2(8-\alpha)}{(5-\alpha)(4-\alpha)(3-\alpha)}$ |
| $B_{6,5}^*$ | $\frac{2(1-\alpha)(8-\alpha)}{(5-\alpha)(4-\alpha)(3-\alpha)}$ |
| $B_{6,6}^*$ | $\frac{2(1-\alpha)(2-\alpha)}{(5-\alpha)(4-\alpha)}$ |
| $B_{7,8}^*$ | $\frac{(1-\alpha)^2(2+\alpha)(10+\alpha)}{(6-\alpha)(5-\alpha)(4-\alpha)(3-\alpha)}$ |
| $B_{7,9}^*$ | $\frac{2(1-\alpha)^2(10+\alpha)}{(6-\alpha)(5-\alpha)(4-\alpha)}$ |
| $B_{7,10}^*$ | $\frac{10(1-\alpha)(2-\alpha)}{(6-\alpha)(5-\alpha)(3-\alpha)}$ |
| $B_{7,11}^*$ | $\frac{5(1-\alpha)^2(2-\alpha)}{(6-\alpha)(5-\alpha)(3-\alpha)}$ |
| $B_{8,13}^*$ | $\frac{8(1-\alpha)^2(1+\alpha)(2+\alpha)(3+\alpha)}{(7-\alpha)(6-\alpha)(5-\alpha)(4-\alpha)(3-\alpha)}$ |
| $B_{8,14}^*$ | $\frac{16(1-\alpha)^2(1+\alpha)(3+\alpha)}{(7-\alpha)(6-\alpha)(5-\alpha)(4-\alpha)}$ |
| $B_{8,15}^*$ | $\frac{8(1-\alpha)^2(3+\alpha)(8-\alpha)}{(7-\alpha)(6-\alpha)(5-\alpha)(4-\alpha)(3-\alpha)}$ |
| $B_{8,16}^*$ | $\frac{4(1-\alpha)^3(3+\alpha)(8-\alpha)}{(7-\alpha)(6-\alpha)(5-\alpha)(4-\alpha)(3-\alpha)}$ |
| $B_{8,17}^*$ | $\frac{8(1-\alpha)^2(2-\alpha)(3+\alpha)}{(7-\alpha)(6-\alpha)(5-\alpha)(4-\alpha)}$ |
| $B_{8,21}^*$ | $\frac{20(1-\alpha)(2-\alpha)}{(7-\alpha)(6-\alpha)(5-\alpha)(3-\alpha)}$ |
| $B_{8,22}^*$ | $\frac{20(1-\alpha)^2(2-\alpha)}{(7-\alpha)(6-\alpha)(5-\alpha)(3-\alpha)}$ |
| $B_{8,23}^*$ | $\frac{5(1-\alpha)^3(2-\alpha)}{(7-\alpha)(6-\alpha)(5-\alpha)(3-\alpha)}$ |

**Table 4:** Probabilities under the β-model of the trees involved in the formula for the variance of *QIB*_n_

| Tree | $P_{\beta,n}^{B,*}$ |
| --- | --- |
| $Q_3^*$ | $\frac{3(\beta+2)}{7\beta+18}$ |
| $B_{5,3}^*$ | $\frac{2(\beta+2)}{3\beta+8}$ |
| $B_{6,4}^*$ | $\frac{45(\beta+2)^2(\beta+4)}{(31\beta^2+194\beta+300)(7\beta+18)}$ |
| $B_{6,5}^*$ | $\frac{60(\beta+2)(\beta+3)(\beta+4)}{(31\beta^2+194\beta+300)(7\beta+18)}$ |
| $B_{6,6}^*$ | $\frac{10(\beta+2)(\beta+3)}{31\beta^2+194\beta+300}$ |
| $B_{7,8}^*$ | $\frac{3(\beta+2)^2(\beta+4)(\beta+5)}{(\beta+3)(3\beta+8)(3\beta+10)(7\beta+18)}$ |
| $B_{7,9}^*$ | $\frac{2(\beta+2)^2(\beta+5)}{(\beta+3)(3\beta+8)(3\beta+10)}$ |
| $B_{7,10}^*$ | $\frac{20(\beta+2)(\beta+3)}{3(3\beta+10)(7\beta+18)}$ |
| $B_{7,11}^*$ | $\frac{5(\beta+2)^2}{(3\beta+10)(7\beta+18)}$ |
| $B_{8,13}^*$ | $\frac{504(\beta+2)^2(\beta+4)^2(\beta+5)^2(\beta+6)}{(127\beta^3+1383\beta^2+4958\beta+5880)(31\beta^2+194\beta+300)(3\beta+8)(7\beta+18)}$ |
| $B_{8,14}^*$ | $\frac{336(\beta+2)^2(\beta+4)(\beta+5)^2(\beta+6)}{(127\beta^3+1383\beta^2+4958\beta+5880)(31\beta^2+194\beta+300)(3\beta+8)}$ |
| $B_{8,15}^*$ | $\frac{1680(\beta+2)^2(\beta+3)(\beta+4)(\beta+5)(\beta+6)}{(127\beta^3+1383\beta^2+4958\beta+5880)(31\beta^2+194\beta+300)(7\beta+18)}$ |
| $B_{8,16}^*$ | $\frac{1260(\beta+2)^3(\beta+4)(\beta+5)(\beta+6)}{(127\beta^3+1383\beta^2+4958\beta+5880)(31\beta^2+194\beta+300)(7\beta+18)}$ |
| $B_{8,17}^*$ | $\frac{280(\beta+2)^2(\beta+3)(\beta+5)(\beta+6)}{(127\beta^3+1383\beta^2+4958\beta+5880)(31\beta^2+194\beta+300)}$ |
| $B_{8,21}^*$ | $\frac{560(\beta+2)(\beta+3)^3(\beta+4)}{(127\beta^3+1383\beta^2+4958\beta+5880)(7\beta+18)^2}$ |
| $B_{8,22}^*$ | $\frac{840(\beta+2)^2(\beta+3)^2(\beta+4)}{(127\beta^3+1383\beta^2+4958\beta+5880)(7\beta+18)^2}$ |
| $B_{8,23}^*$ | $\frac{315(\beta+2)^3(\beta+3)(\beta+4)}{(127\beta^3+1383\beta^2+4958\beta+5880)(7\beta+18)^2}$ |

